# An Epigenomic Roadmap Primes Non-Growing Oocytes for Maturation and Early Embryogenesis

**DOI:** 10.1101/2025.07.06.663377

**Authors:** Mengwen Hu, Yasuhisa Munakata, Yu-Han Yeh, Han Wang, Neil Hunter, Richard M. Schultz, Satoshi H. Namekawa

## Abstract

Female reproductive lifespan is defined by long-lived, non-growing oocytes (NGOs) that comprise the ovarian reserve. NGOs are assumed to acquire the epigenetic marks that will define the early embryo only after they exit the ovarian reserve and become activated for growth. Contrary to this dogma, we show that mouse NGOs possess abundant histone modifications that both underlie maintenance of the ovarian reserve and prime the epigenome of growing oocytes for early embryogenesis. As NGOs are established around birth, Polycomb Repressive Complex 1 (PRC1) mediates abundant H2AK119 ubiquitylation and reprograms the H3K27 acetylation landscape, which is essential to maintain the ovarian reserve. Importantly, the PRC1-driven epigenetic state of NGOs provides a blueprint for subsequent generation of a PRC2-catalyzed H3K27 trimethylation profile in growing oocytes that is characterized by broad domains and DNA methylation-independent imprints that are transmitted to the embryo. Thus, Polycomb complexes play pivotal roles in priming the NGO epigenome for oocyte maturation and early embryogenesis.

**Highlights:** Non-growing oocytes have rich epigenetic landscapes to prime subsequent development Reprogramming of H3K27ac by PRC1 is required to maintain non-growing oocytes PRC1-catalyzed H2AK119ub provides a blueprint for PRC2-H3K27me3 in growing oocytes In non-growing oocytes, PRC1 primes DNA methylation-independent H3K27me3-imprinting

## Introduction

Oogenesis begins in the embryo where female primordial germ cells (PGCs) rapidly divide and form germline cysts of inter-connected oogonia through incomplete cytokinesis. Following epigenetic reprogramming, including genome-wide DNA demethylation, female PGCs enter meiosis^1^. During meiotic prophase I (MPI), homologous chromosomes pair, synapse, recombine, and then desynapse before oocytes arrest in the dictyate stage around birth. As meiosis arrests, germline cysts break down into single non-growing oocytes (NGOs) surrounded by a layer of flattened pre-granulosa cells to form primordial follicles^2^. Approximately two-thirds of the oocytes die during these transitions^3^. The resulting pool of primordial follicles constitutes the ovarian reserve that determines female reproductive lifespan, sustaining fertility for months in mice and for decades in humans^3–5^.

Progressive depletion of the ovarian reserve occurs with each estrous cycle as cohorts of NGOs are recruited for growth and maturation. As growing oocytes (GOs) enlarge, they accumulate essential proteins, RNAs, and organelles. Dominant follicles continue to mature into full-grown oocytes (FGOs) while the other GOs cease growing and die^6^. Following a gonadotropin surge, FGOs complete meiosis I, arrest again at metaphase II, are ovulated, and await fertilization, which triggers completion of meiosis and formation of a zygote^7^.

Following epigenetic reprogramming in PGCs, the oocyte genome remains hypomethylated until oocyte growth, during which de novo DNA methylation occurs, and a variety of histone modifications are established^8–11^. Previous analysis suggests that DNA hypomethylation in NGOs is accompanied by scarce epigenetic marks^12^. Therefore, if and how chromatin-based mechanisms enable the establishment and maintenance of the ovarian reserve remains unclear. This knowledge gap persists, at least in part, because of the difficulty in isolating sufficient numbers of NGOs for epigenomic analyses^13^.

We previously showed that Polycomb Repressive Complex 1 (PRC1) plays an essential role in establishing the ovarian reserve by repressing the MPI transcriptional program^14^. Polycomb group proteins (PcGs) consist of two major, functionally related complexes—PRC1 and PRC2, which catalyze two major repressive histone marks involved in the repression of non-lineage-specific genes and define cellular identity during development^15–17^. PRC1 monoubiquitinates histone H2A at lysine 119 (H2AK119ub), and PRC2 trimethylates H3 at lysine 27 (H3K27me3). With respect to the formation and maintenance of the ovarian reserve, the spatiotemporal dynamics, functions, and interplay between H2AK119ub and H3K27me3 remain unknown.

Mouse FGOs exhibit unique ‘noncanonical’ landscapes of histone modifications. For example, H3K4me3, an active mark that is typically enriched at promoters, is present in broad domains that are maintained in early embryos prior to zygotic genome activation^18,19^ (ZGA). Another active mark, H3K27 acetylation (H3K27ac), typically enriched at both promoters and active enhancers, becomes prevalent in gene deserts of the FGO genome^11^. Repressive H3K27me3 is widely distributed throughout the FGO genome, covering most regions that are not actively transcribed^20^; a pattern that persists until the blastocyst stage and mediates DNA methylation-independent imprinting that regulates paternal allele-specific gene expression and imprinted X-chromosome inactivation in early embryos^21–23^. H2AK119ub largely coexists with H3K27me3 in FGOs, but their distribution becomes distinct after fertilization^24,25^.

How the NGO epigenome is established and maintained during arrest, whether it is reprogrammed in GOs, and how it relates to the FGO epigenome are key questions that remain largely unanswered. Here, we present a comprehensive and quantitative epigenomic roadmap of mouse perinatal oocytes and show that Polycomb-based mechanisms govern epigenetic reprogramming during these stages. Together, our data reveal that epigenetic priming in NGOs shapes the epigenetic landscape during oocyte growth and the oocyte-to-embryo transition, and H3K27me3-mediated imprinting can be traced all the way back to the establishment of non-growing oocytes around the time of birth and is regulated by PRC1-mediated H2AK119ub.

## Results

### Two major epigenomic transitions occur during perinatal oogenesis

In mice, critical events associated with formation of the ovarian reserve occur around birth and include the breakdown of germline cysts, dictyate arrest, and assembly of primordial follicles (Figure 1A)^26–29^. Concurrently, perinatal oocytes undergo the Perinatal Oocyte Transition (POT), exiting meiotic prophase I (MPI) to become NGOs^14,30^. A second major transition, the Primordial-to-Primary follicle Transition (PPT), occurs when NGOs are recruited to initiate growth^31^. Both transitions are associated with dramatic transcriptional changes in the oocyte^30,32^. To determine the epigenomic landscapes that underlie the PPT and POT, we profiled the genomic distribution of four key histone modifications, H2AK119ub, H3K27me3, H3K4me3, and H3K27ac in wild-type (WT) perinatal oocytes at four developmental stages: MPI oocytes at E18.5 (E18O), oocytes transitioning to dictyate arrest at postnatal day one (P1 oocytes, P1O), NGOs within primordial follicles at P5 to 6 (NGO) and early GOs at P6 to 7 (GO) (Figures 1B, C). To quantify changes between stages, we employed low-input cleavage under targets and tagmentation (CUT&Tag)^33,34^ with spike-in controls normalized for cell numbers. After confirming the high quality and reproducibility of CUT&Tag datasets (Figure S1A), biological replicates were pooled for downstream analyses.

**Figure 1.**
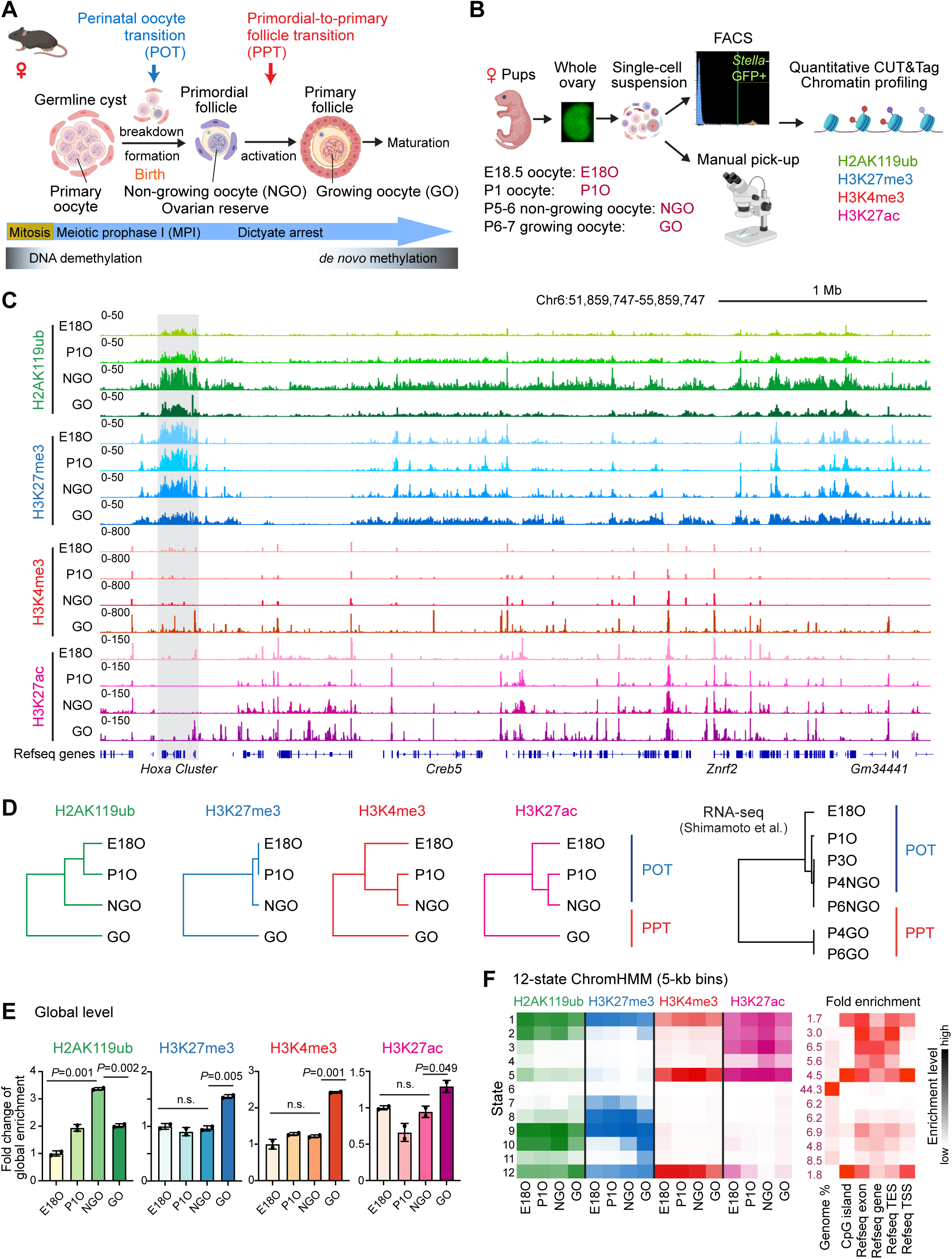
Dynamic epigenomic landscapes in perinatal oocytes. (A) Schematic of mouse perinatal oogenesis (created with BioRender.com). DNA methylation levels are represented with a graph. NGO, non-growing oocyte; GO, growing oocyte. (B) Overview of the experimental steps for chromatin profiling using quantitative CUT& Tag on perinatal oocytes (created with BioRender.com). (C) Genome browser views of H2AK119ub, H3K27me3, H3K4me3 and H3K27ac landscapes during perinatal oogenesis. (D) Hierarchical clustering of global H2AK119ub, H3K27me3, H3K4me3, and H3K27ac in 10-kb bins and hierarchical clustering of gene expression across all stages during perinatal oogenesis in wild-type. E18O represents oocytes in meiotic prophase I; P1O and P3O represent oocytes transitioning to dictyate arrest; P4 and P6 small oocytes represent NGO residing in primordial follicles; and P4 and P6 large oocytes represent GO in primary follicles after initiation of oocyte growth. (E) Bar chart showing the foldchange of global levels of H2AK119ub, H3K27me3, H3K4me3, and H3K27ac as determined by quantitative CUT&Tag in perinatal oocytes (n = 2 biological replicates, indicated by dots). P values of pairwise comparisons (two-sided unpaired Student’s t-test) are given. n.s., not significant. Data are represented as mean ± SD. (F) Heatmap illustrating chromatin state dynamics in perinatal oocytes using ChromHMM analysis. The intensity of colors indicates the enrichment level for each modification belonging to the given state at each stage. Genome distributions of all 12 states were shown on the right. The intensity of colors indicates the fold enrichment level of the given state. The number in the first column represents the genome coverage percentage of each state.

Hierarchical clustering revealed two major epigenomic transitions corresponding to the POT and PPT (Figure 1D), which were mirrored by dynamic changes in the oocyte transcriptome (Figure 1D, right). During POT, the global H2AK119ub progressively increased (∼3.4-fold, Figures 1E and S1B), accompanied by increases in peak width and intensity (Figures S2A-C). In contrast, H3K27me3 and H3K4me3 levels remained relatively constant (Figures 1E and S1B), with similar peak numbers, widths, and intensities (Figures S2E-G, S3A-C). Although global H3K27ac levels did not change significantly during POT (Figure 1E), peak numbers progressively increased between E18Os to NGOs (Figure S3E-G). Subsequently, upon PPT, H2AK119ub levels decreased while H3K27me3, H3K4me3, and H3K27ac increased (Figures 1E and S1B).

Patterning of the four histone marks was also dynamic during POT and PPT. For example, extensive gain of H3K4me3 during PPT and progressive gain of H3K27ac were observed (Figures S3A, E). Changes in genomic distributions were analyzed using ChromHMM^35^, which classifies regions into 12 chromatin states, identifying dynamic, region-specific epigenomic changes during perinatal oogenesis (Figure 1F). States 1, 5, and 12 were characterized by high H3K4me3 and H3K27ac levels and enrichment for CpG islands (CGIs), representing putative gene promoters. Among them, States 1 and 12 were also enriched for repressive H2AK119ub and H3K27me3, representing putative bivalent promoters. Strikingly, ChromHMM identified genome-wide redistribution of H3K27ac during the POT.

During PPT, dynamic redistribution of Polycomb marks was detected during the PPT that suggests a mechanistic link between PRC1-mediated H2AK119ub in NGOs and ensuing PRC2-mediated H3K27me3 in GOs. In GOs, ChromHMM States 2, 10, and 11 showed a gain of H3K27me3. Prior to this gain, in NGOs, the ChromHMM states for H2AK119ub and H3K27me3 were distinct but then in GOs became much more similar (Figure 1F). Consistently, the genomic distribution of H3K27me3 in GOs became similar to that of H2AK119ub in both NGOs and GOs (Figure 1C). Gain of H3K27me3 in GOs occurred in broad regions that are distinct from representative Polycomb target loci, such as the *Hoxa* gene cluster (Figure 1C, gray box), suggesting that H3K27me3 is a unique regulated in GOs.

The two major shifts in chromatin states during the POT and PPT were also associated with changes in expression of pertinent chromatin-modifying enzymes related to the four histone modifications (Figure S4). Taken together, this analysis reveals dynamic epigenomic changes during the POT, as the ovarian reserve is established, and during the PPT as oocytes initiate growth.

### Coordinated chromatin changes drive transcriptional transitions during the POT and PPT

Global analysis of annotated gene promoters confirmed local loss of H3K27ac and gain of H2AK119ub in NGOs and subsequent gain of H3K27me3 in GOs (Figure S5A). In mammals, a large group of promoters contains CGIs that are generally constitutively unmethylated and enriched for PRC1 and PRC2 and their associated marks H2AK119ub and H3K27me3, respectively^36,37^. Comparison of histone marks at 14,040 CGI-associated genes (CGI genes) and 39,755 genes without CGIs (non-CGI genes) revealed extensive reprogramming of CGI gene promoters (Figure 2A). A gain of H2AK119ub was observed at ∼73% (n=10,250) of the transcription start sites (TSSs) of CGI gene during the POT (Figure 2B), peaking in NGOs (Figure 2A). H3K27me3 enrichment followed the H2AK119ub pattern, peaking in GOs (Figure 2A). Conversely, a sharp loss of H3K27ac was observed at ∼62% (n=8,615) of CGI gene TSSs during the POT (Figure 2B), whereas H3K4me3 levels were relatively constant (Figure 2A). These changes were unexpected because genome-wide H3K27ac often coexisted with H3K4me3 (Figure S3I). Subsequently, during the PPT, the active marks H3K4me3 and H3K27ac were increased, reflecting the global gene activation that occurs in GOs.

**Figure 2.**
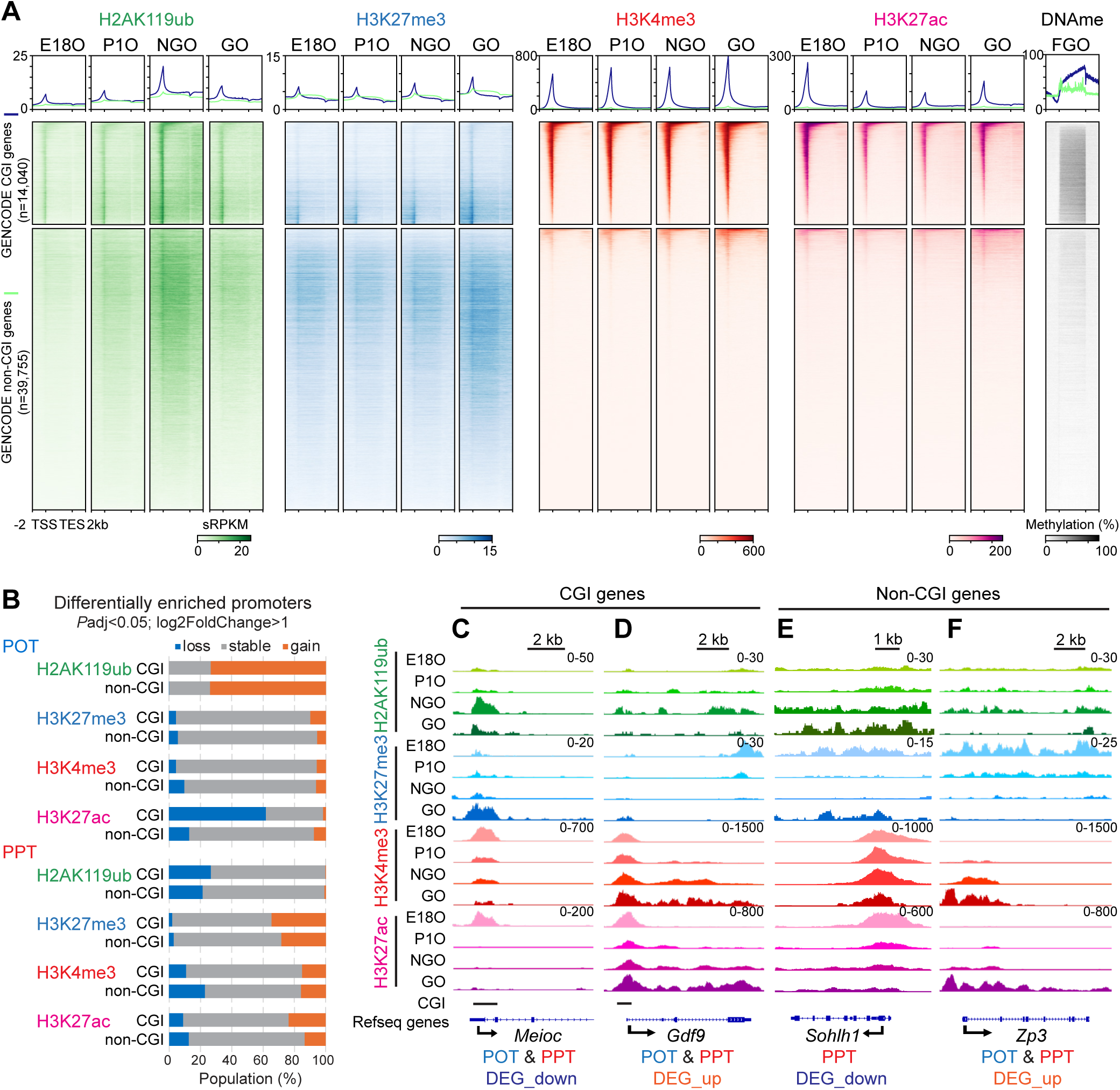
Coordinated chromatin changes drive transcriptional transitions during POT and PPT. (A) Average tag density plots and heatmaps showing H2AK119ub, H3K27me3, H3K4me3, and H3K27ac dynamics at gene body regions (TSS -TES ± 2 kb) of CGI genes and non-CGI genes in perinatal oocytes. The color keys represent signal intensity, and the numbers represent spike-in scaled RPKM (sRPKM) values. Right: average tag density plot and heatmap showing DNA methylation levels in full-grown oocytes (FGO) of corresponding CGI and non-CGI gene groups. The color keys represent signal intensity, and the numbers represent DNA methylation percentage. (B) Bar charts showing percentages of differentially enriched promoters of CGI and non-CGI genes for H2AK119ub, H3K27me3, H3K4me3, and H3K27ac during POT and PPT. (C-F) Track views of the representative POT or PPT DEGs in CGI gene group (*Meioc* and *Gdf9*) and non-CGI gene group (*Sohlh1* and *Zp3*) gene loci showing differential H2AK119ub, H3K27me3, H3K4me3, and H3K27ac occupancy during perinatal oogenesis. Data ranges in the upper right represent sRPKM values from combined replicates.

To further explore the relationship between chromatin state and gene regulation during the POT and PPT, we analyzed genes that were differently expressed (DEGs) during these transitions (identified from publicly available data^30^, Figure S5B). *Meioc* is a representative CGI gene and a critical regulator of MPI^38^, that is expressed in MPI and suppressed during the POT and PPT (Figure S5B). At the *Meioc* promoter, loss of H3K27ac and gain of H2AK119ub was observed during the POT, followed by gain of H3K27me3 in GOs (Figure 2C). *Gdf9* is also a CGI gene and an essential regulator of folliculogenesis^39^ whose expression is up-regulated during the POT and PPT (Figure S5B). During the POT, loss of H3K27ac and gain of H2AK119ub were observed at the *Gdf9* promoter while H3K4me3 and H3K27ac increased in the gene body as it was expressed (Figure 2D). Genome-wide analysis further confirmed that global loss of H3K27ac and gain of H2AK119ub at promotes during the POT were frequently associated with both up- and down-regulated genes (Figure S5C). This epigenetic signature suggests a relatively repressive chromatin state at promoters after the POT, which could be critical for maintaining long-term stability of the ovarian reserve.

In GOs, transcription-dependent DNA methylation is gradually established^40^. We reanalyzed previous whole-genome bisulfite sequencing data of FGOs^8^ and found that at CGI genes, a global gain of DNA methylation occurs in gene bodies while the TSSs remained unmethylated (Figure 2A). Thus, the reprogramming of CGI-gene promoters (loss of H3K27ac and gain of H2AK119ub) during the POT and subsequent gain of H3K27me3 during the PPT underlie protection from *de novo* DNA methylation in GOs and shape the DNA methylation landscape in FGOs.

In the gene bodies of non-CGI genes, a global gain of H2AK119ub was seen in NGOs, followed by a gain of H3K27me3 in GOs (Figure 2A). *Sohlh1* is a representative non-CGI gene required for oogenesis^41^, that is expressed in NGOs and downregulated during the PPT. Unexpectedly, when *Sohlh1* was expressed in NGOs, the gene body was covered with H2AK119ub. Subsequent down-regulation of *Sohlh1* in GOs was associated with a further gain of H2AK119ub and increased H3K27me3 (Figure 2E). Another non-CGI gene, *Zp3*, encoding the Zona pellucida 3 (ZP3) protein that is essential for fertility ^42,43^, is expressed in both NGOs and GOs. Similar to *Sohlh1,* H2AK119ub covered the gene body when *Zp3* was expressed in NGOs (Figure 2F). However, *Zp3* induction in GOs was associated with reduced H2AK119ub and H3K27me3 and increased H3K4me3 and H3K27ac. Thus, in contrast to the promoter reprogramming of CGI genes, non-CGI genes undergo distinct global epigenetic programming within the gene bodies.

### PRC1 regulates H3K27ac reprogramming during ovarian reserve formation

Because H3K27ac reprogramming is a key feature of POT, we further examined the epigenomic features at H3K27ac peaks during POT. 17,481 peaks lost H3K27ac intensity, and these peaks were highly enriched with H3K4me3, as well as the Polycomb marks H2AK119ub and H3K27me3 (“POT loss” peaks: Figure 3A). These “POT loss” peaks substantially gained H2AK119ub in NGOs (Figure 3A). Conversely, 33,667 newly established H3K27ac peaks in NGOs modestly gained H2AK119ub but not H3K4me3 and H3K27me3 (“POT gain” peaks), and the 62,127 unchanged H3K27ac peaks also modestly gained H2AK119ub in NGOs (“POT stable” peaks). Overall, these data suggest that Polycomb is involved in H3K27ac reprogramming during POT. Consistent with this possibility, PRC2-H3K27me3 is mutually exclusive with H3K27ac because they are at the same residue, and PRC1 potentially modulates H3K27ac through interactions with a histone acetylase^44^ or deacetylase^45^.

**Figure 3.**
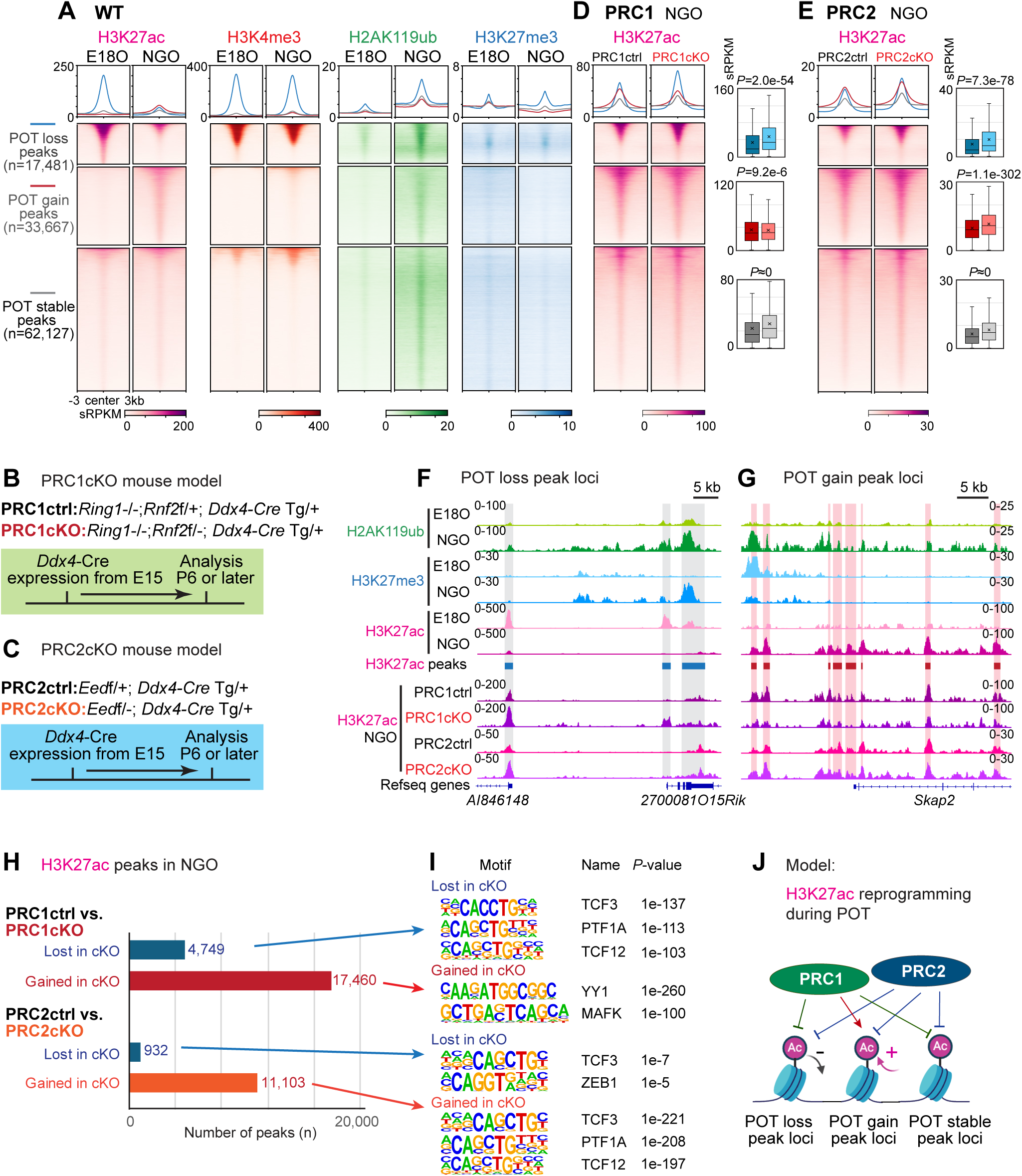
PRC1 regulates H3K27ac reprogramming during ovarian reserve formation. (A, D and E) Average tag density plots and heatmaps showing histone modification changes at differential H3K27ac peaks region (peak center ± 3 kb) in E18O and NGO of WT (A), NGO of PRC1ctr&cKO (D), NGO of PRC2ctrl&cKO (E), respectively. Right: Box and whisker plots showing quantifications of H3K27ac enrichment within peaks (sRPKM values) of indicated peak groups in PRC1ctr&cKO, PRC2ctrl&cKO, respectively. Boxes show the 25th and 75th percentile with the median, and whiskers indicate 1.5 times the interquartile range. *P* values of pairwise comparisons (two-sided unpaired Student’s t-test) are given. (B and C) Schematic of mouse models and experiments. (F and G) Track views of the representative POT loss (F) or POT gain (G) H3K27ac peaks region showing H2AK119ub, H3K27me3, and H3K27ac occupancy during POT in WT, and H3K27ac changes in PRC1ctr&cKO, PRC2ctrl&cKO NGO. Data ranges represent sRPKM values from combined replicates. (H and I) Bar chart showing numbers (H) and motif analysis (I) of differentially enriched H3K27ac peaks in NGO of PRC1ctrl&cKO, PRC2ctrl&cKO. (J) A schematic model showing the regulation of H3K27ac reprogramming during POT by Polycomb.

To determine the role of Polycomb complexes in H3K27ac reprogramming, we used two germ cell-specific conditional knockout mouse models that lead to loss of function of PRC1 or PRC2. Our previously established PRC1 loss-of-function model (PRC1cKO)^14,46,47^ uses *Ddx4*-Cre, which is expressed in germ cells from E15^48^, to inactivate the E3 ubiquitin ligase RNF2 in a genetic background that lacks the partially redundant paralog, RING1(RING1A)^49,50^ (Figure 3B). We also generated a PRC2 loss-of-function model (termed PRC2cKO) by crossing an *Eed* (an essential component of PRC2) floxed mouse line^51^ with a *Ddx4*-Cre expressing line to prevent H3K27me3 deposition from E15 (Figure 3C). Quantitative CUT&Tag analyses confirmed that PRC2cKO resulted in markedly decreased H3K27me3 in NGOs (Figures S6A-E).

Notably, PRC1 regulates H3K27ac in a site-specific manner. Although the global H3K27ac levels were generally higher in PRC1cKO NGOs compared to PRC1ctrl NGOs (Figures S7A-D), the H3K27ac levels remained particularly high at the “POT loss” peak loci in PRC1cKO NGOs, whereas the H3K27ac levels were low at the “POT gain” peak loci in PRC1cKO NGOs (Figure 3D). In contrast, in PRC2cKO NGOs, the H3K27ac levels were higher across all groups, presumably reflecting the antagonistic relationship between H3K27me3 and H3K27ac (Figure 3E). In WT during POT, there was a gain of H2AK119ub at “POT loss” peaks (Figure 3F) and at “POT gain” peaks (Figure 3G), and PRC1 facilitates this loss and gain of H3K27ac during POT (Figures 3F, G). In PRC1cKO NGOs, 17,460 H3K27ac peaks were differentially gained compared to PRC1ctrl NGOs, and these peaks were enriched for motifs of YY1 and MAFK (Figures 3H, I). These TF motifs were also found in the “POT loss” peaks (Figure S3J), supporting the notion that PRC1 facilitates H3K27ac peak loss during POT in WT. Conversely, the 4,749 H3K27ac peaks were differentially lost compared to PRC1ctrl NGOs, and these peaks were enriched for motifs of TCF3 (E2A), PTF1A, and TCF13 (HEB), which are identified in the “POT gain” peaks in WT oocytes (Figure S3J), further supporting the notion that PRC1 also facilitates H3K27ac gain during POT. Importantly, TCF3/12 are critical regulators of postnatal oogenesis/ folliculogenesis^11^. In contrast, in PRC2cKO NGOs, both the gained and lost H3K27ac peaks were similarly enriched for TCF3/12 motifs (Figure 3I). Thus, PRC2 is not involved in the developmental transition during POT but only restricts the H3K27ac levels. The contrasting effects between PRC1cKO and PRC2cKO reveal distinct functions for PRC1 and PRC2: PRC1 facilitates both H3K27ac loss and gain during POT, whereas PRC2-mediated H3K27me3 generally antagonizes H3K27ac (Figure 3J). Taken together, these analyses identified PRC1 as a key regulator of the H3K27ac reprogramming at gene regulatory elements during ovarian reserve formation.

### Distinct functions of PRC1 and PRC2 in gene repression in non-growing oocytes

Because we found distinct functions of PRC1 and PRC2 in H3K27ac reprogramming during POT, we next sought to clarify the functional relationship between PRC1 and PRC2 in NGOs. Quantitative CUT&Tag analysis revealed that, in PRC1cKO NGOs, H3K27me3 levels and distribution remained largely unchanged (Figures 4A and S8A). The same was true for H2AK119ub in PRC2cKO NGOs, which showed only a modest decrease in total enrichment level (Figures 4B and S8B). Thus, PRC1 and PRC2 appear to function largely independent of each other in NGOs.

**Figure 4.**
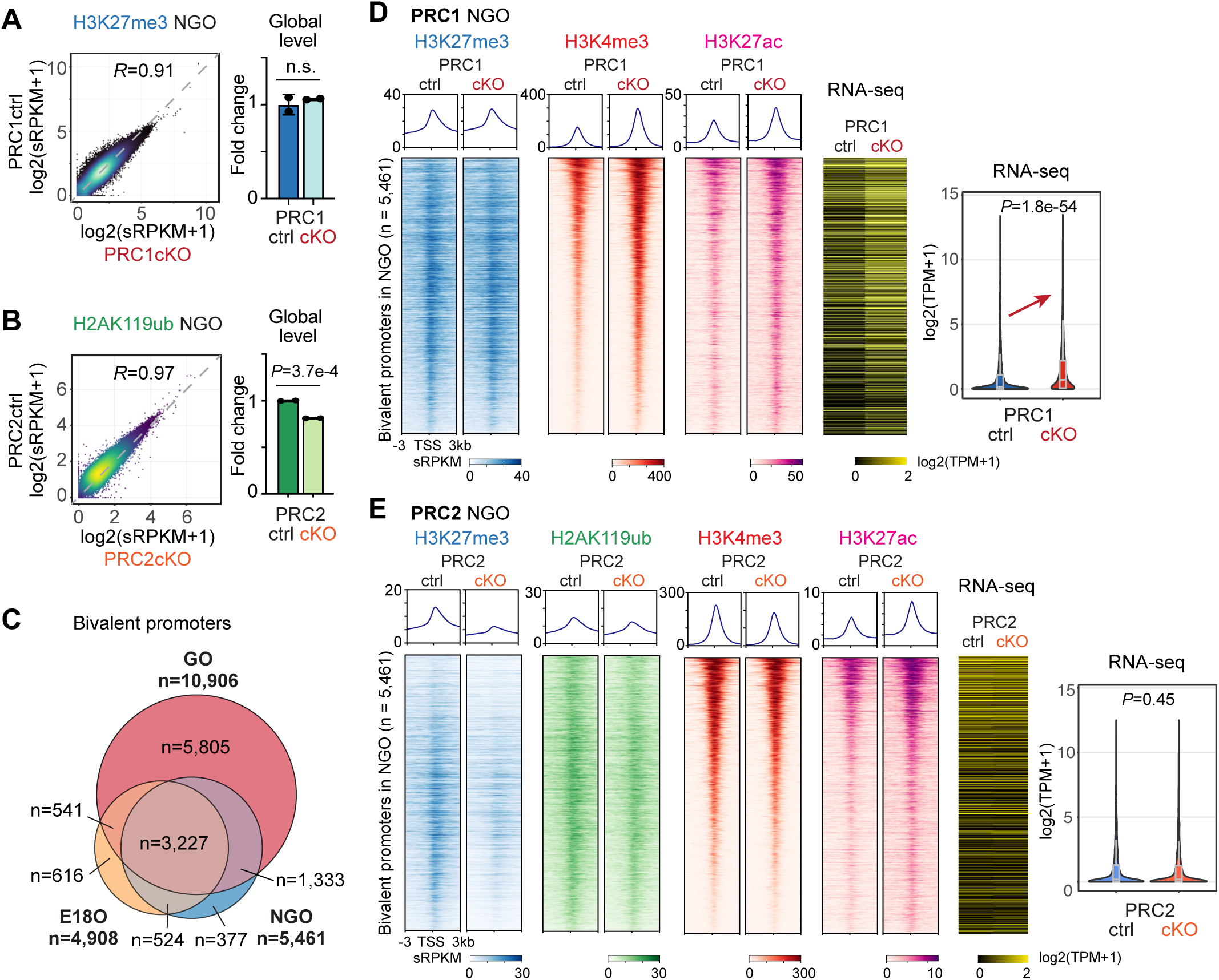
Distinct functions of PRC1 and PRC2 in gene repression in non-growing oocytes. (A and B) Density scatter plots showing genome-wide H3K27me3 enrichment by 10-kb bins comparing PRC1ctrl and PRC1cKO NGO (A), or H2AK119ub in PRC2ctrl and PRC2cKO NGO (B). The Pearson correlation coefficient is indicated. The bar charts on the right show the foldchange of H3K27me3 global level in PRC1ctrl and PRC1cKO NGO (A) or H2AK119ub in PRC2ctrl and PRC2cKO NGO (B). (n = 2 biological replicates, indicated by dots). *P* values of pairwise comparisons (two-sided unpaired Student’s t-test) are given. n.s., not significant. Data are represented as mean ± SD. (C) Venn diagrams showing bivalent promoters dynamics in perinatal oocytes. Numbers and percentages of total bivalent promoters are indicated. (D and E) Average tag density plots and heatmaps showing histone modifications changes at WT NGO bivalent promoter regions (TSS ± 3 kb) in PRC1ctrl&cKO NGO (D) or PRC2ctrl&cKO NGO (E). The color keys represent signal intensity, and the numbers represent sRPKM values. Right: Heat map and violin plots with included boxplots showing RNA expression (log2-transformed TPM) of corresponding bivalent genes in PRC1ctrl&cKO NGO (D) or PRC2ctrl&cKO NGO (E). Boxes show the 25th and 75th percentile with the median, and whiskers indicate 1.5 times the interquartile range. *P* values of pairwise comparisons (two-sided unpaired Student’s t-test) are given.

Next, we examined if the two Polycomb complexes regulate bivalent promoters, which are characterized by the presence of both H3K4me3 and H3K27me3 and are typically poised for later activation^52^. Promoter bivalency plays a critical role in development and differentiation, particularly in the context of the germline^53,54^. We identified 4,908 and 5,461 bivalent promoters in E18Os and NGOs, respectively, with substantial overlap (3,751 in common), indicating an overall consistent feature of promoter bivalency during POT (Figure 4C). The number of bivalent promoters increased substantially upon PPT (10,906 in GOs) due to the widespread increase of H3K27me3 (Figures 2A, 4C, and S5A). As a key characteristic of promoter bivalency, we profiled H3K4me3 in both PRC1cKO and PRC2cKO NGOs by quantitative CUT&Tag (Figures S9A, B). Unexpectedly, the effects of PRC1 and PRC2 deficiency on H3K4me3 were opposite: PRC1 depletion significantly increased global H3K4me3 levels, especially at peaks and promoter regions (Figures S9C-F), whereas PRC2 deletion resulted in a modest genome-wide decrease in H3K4me3 levels (Figures S9A-F). At bivalent promoters, PRC1 depletion resulted in increased H3K4me3 enrichment, whereas H3K27me3 remained unchanged, leading to ectopic expression of bivalent genes in NGOs (Figure 4D). Promoters that gained H3K4me3 in PRC1cKO NGOs largely overlapped with Polycomb-binding sites identified in many other cell types (Figure S9G), consistent with H2AK119ub counteracting H3K4me3 at promoters^17,55^. In contrast, at bivalent promoters, H3K27me3 coverage was depleted as expected after PRC2 deletion, whereas the H2AK119ub profile remained unchanged, and H3K4me3 levels were modestly reduced, resulting in no significant change in expression of bivalent genes (Figure 4E). Consistent with this observation, only a modest change in gene expression was observed in PRC2cKO NGOs (Figures S10A, B), compared to a substantial change in gene expression in PRC1cKO NGOs^14^. These results suggest that in NGOs, H3K27me3 and H3K4me3 do not have an antagonistic relationship as observed in ESCs^56^, and that PRC1 plays a major role in the regulation of bivalent promoters in NGOs.

### PRC1-H2AK119ub in the ovarian reserve programs PRC2-H3K27me3 in growing oocytes

Next, we sought to determine the mechanism underlying the epigenomic transition during PPT. Based on the similarity between the H2AK119ub pattern in NGOs and the H3K27me3 pattern in GOs (Figure 1C, F), we investigated the interplay between PRC1-H2AK119ub and PRC2-H3K27me3. Hierarchical clustering analysis of our data with previously published Polycomb CUT&RUN data in FGOs^24^ revealed three major clusters reflecting sequential transitions (Figure 5A): H3K27me3 in small oocytes before growth (Cluster I); H2AK119ub in small oocytes and H3K27me3 in GOs (Cluster II); and H2AK119ub in GOs and both marks in FGOs (Cluster III). Remarkably, compared to H3K27me3 in earlier stages, the global distribution of H3K27me3 in GOs was more highly correlated with H2AK119ub in NGOs (Pearson correlation=0.87) (Figures 5A, B). Furthermore, H2AK119ub peaks in NGOs and H3K27me3 peaks in GOs largely overlapped with each other (Figures S2I, J). These results raised the possibility that H2AK119ub in NGOs serves as a blueprint for global reprogramming of H3K27me3 upon PPT.

**Figure 5.**
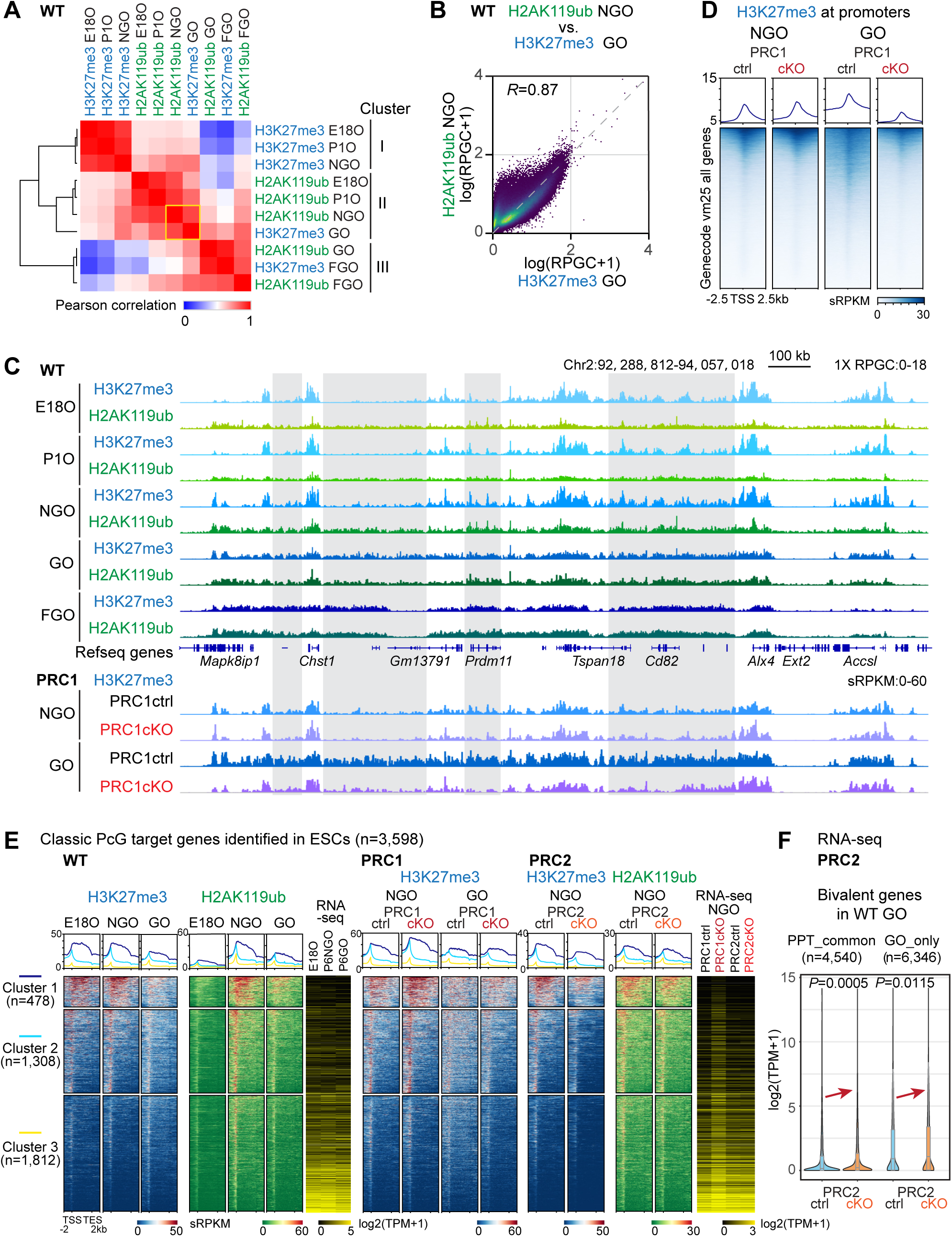
PRC1-H2AK119ub in the ovarian reserve programs PRC2-H3K27me3 in growing oocytes. (A) Heatmaps with hierarchical clustering showing the Pearson correlation of PRC1-H2AK119ub and PRC2-H3K27me3 genome-wide distribution (Reads per genomic content :RPGC) in different oocyte developmental stages. (B) Density scatter plots showing genome-wide distribution between H2AK119ub in WT NGO and H3K27me3 in WT GO by 10-kb bins. The Pearson correlation coefficient is indicated. (C) Genome browser views of H2AK119ub and H3K27me3 landscapes in different oocyte development stages of WT, and H3K27me3 in PRC1ctrl&cKO oocytes. (D) Average tag density plots and heatmaps showing H3K27me3 changes at promoter regions (TSS ± 2.5 kb) in PRC1ctrl and PRC1cKO oocytes. The color keys represent signal intensity, and the numbers represent sRPKM values. (E) Average tag density plots and heatmaps of k-means clusters showing H2AK119ub and H3K27me3 enrichment at gene regions (TSS-TES ± 2 kb) of classic Polycomb-target genes in WT, PRC1, and PRC2 oocytes of indicated genotypes. Right: heatmaps showing RNA-seq expression (log2-transformed TPM) of corresponding k-means clusters in WT, PRC1, and PRC2 oocytes of indicated genotypes. (F) Violin plots with included boxplots showing RNA expression (log2-transformed TPM) of WT GO bivalent genes in PRC2ctrl&cKO GO. Boxes show the 25th and 75th percentile with the median, and whiskers indicate 1.5 times the interquartile range. *P* values of pairwise comparisons (two-sided unpaired Student’s t-test) are given.

Despite the challenge of isolating rare GOs from PRC1cKO due to an early oocyte depletion phenotype^14^, we collected a sufficient number of GOs to determine whether H2AK119ub in NGOs is critical for H3K27me3 reprogramming upon PPT. Remarkably, the H3K27me3 landscape in PRC1cKO GOs remained highly similar to H3K27me3 in NGOs of both PRC1ctrl and PRC2cKO (Figures 5C, S6A and 8C-E). This feature was particularly evident at regions outside promoters, where H3K27me3 normally exhibited a broad distribution similar to H2AK119ub in NGOs upon initiation of oocyte growth. In PRC1cKO GOs, however, we observed a canonical pattern (enrichment at developmental gene promoters) (Figure 5C). Based on these data, we conclude that H2AK119ub in NGOs serves as a blueprint to direct the reprogramming of H3K27me3 during PPT.

To distinguish the feature of the noncanonical distribution of H3K27me3 from its typical distribution at classical Polycomb target genes, we examined the distribution of H3K27me3 and H2AK119ub across the classical Polycomb target genes (3,598 genes) previously identified in ESCs^20^. Clustering analysis categorized these genes into three major groups based on the level of enrichment (Figure 5E). Cluster 1 genes showed the highest levels of both marks throughout the gene bodies, Cluster 2 showed higher enrichment at promoters, and Cluster 3 showed weak enrichment overall. Enrichment levels of Polycomb marks were inversely correlated with transcription, confirming their repressive functions. Whereas H2AK119ub distribution and gene expression were unchanged in PRC2cKO NGOs, PRC1 depletion resulted in substantial up-regulation of Polycomb target genes and, unexpectedly, despite a slight increase in H3K27me3 enrichment at Cluster I genes (Figure 5E), confirming a pivotal role of PRC1 in gene repression in NGOs. Notably, Class I genes maintained H3K27me3 enrichment in GOs after PRC1 depletion, suggesting that PRC1-directed H3K27me3 increase occurs specifically outside of the classical Polycomb target genes, further confirming the noncanonical nature of H3K27me3 reprogramming in PPT.

Concurrent with the extensive gain of H3K27me3 in GOs, numerous bivalent promoters were newly established during PPT (Figure 4C). Although depletion of PRC2-H3K27me3 did not alter expression of genes with bivalent promoters in NGOs (Figure 4e), in GOs, the bivalent genes tended to be up-regulated upon H3K27me3 loss, including the genes common to NGOs and GOs as well as for newly formed bivalent promoter genes in GOs (Figure 5F). These results indicate that PRC2 begins to play a role in gene repression as oocyte growth commences.

### Broad Polycomb domains and H3K27me3-dependent imprinting are primed in non-growing oocytes

A prominent feature of FGO is the presence of noncanonical, distal-rich, broad H3K27me3 and H2AK119ub domains^20,24,25^. To determine when and how these domains emerge, we analyzed enrichment of these marks in perinatal oocytes at genomic loci of all broad peaks (>10 kb) identified in FGOs (Figures 6A, B). We found that broad H3K27me3 domains were already present in GOs (Figure 6A). Notably, the boundaries of these domains were delineated in NGOs (Figure 6A), suggesting that at the onset of oocyte growth, the spreading of H3K27me3 between boundaries leads to formation of the broad domains. This spreading depended on H2AK119ub because it did not occur in PRC1cKO GOs (Figure 6A). Broad H2AK119ub domains could be traced back to NGOs prior to growth (Figure 6B). The H2AK119ub distribution pattern persisted into GOs, despite a decrease in enrichment level. This H2AK119ub pattern was not altered by PRC2 depletion (Figure 6B), confirming that H3K27me3 is largely dispensable for global H2AK119ub modification in NGOs.

**Figure 6.**
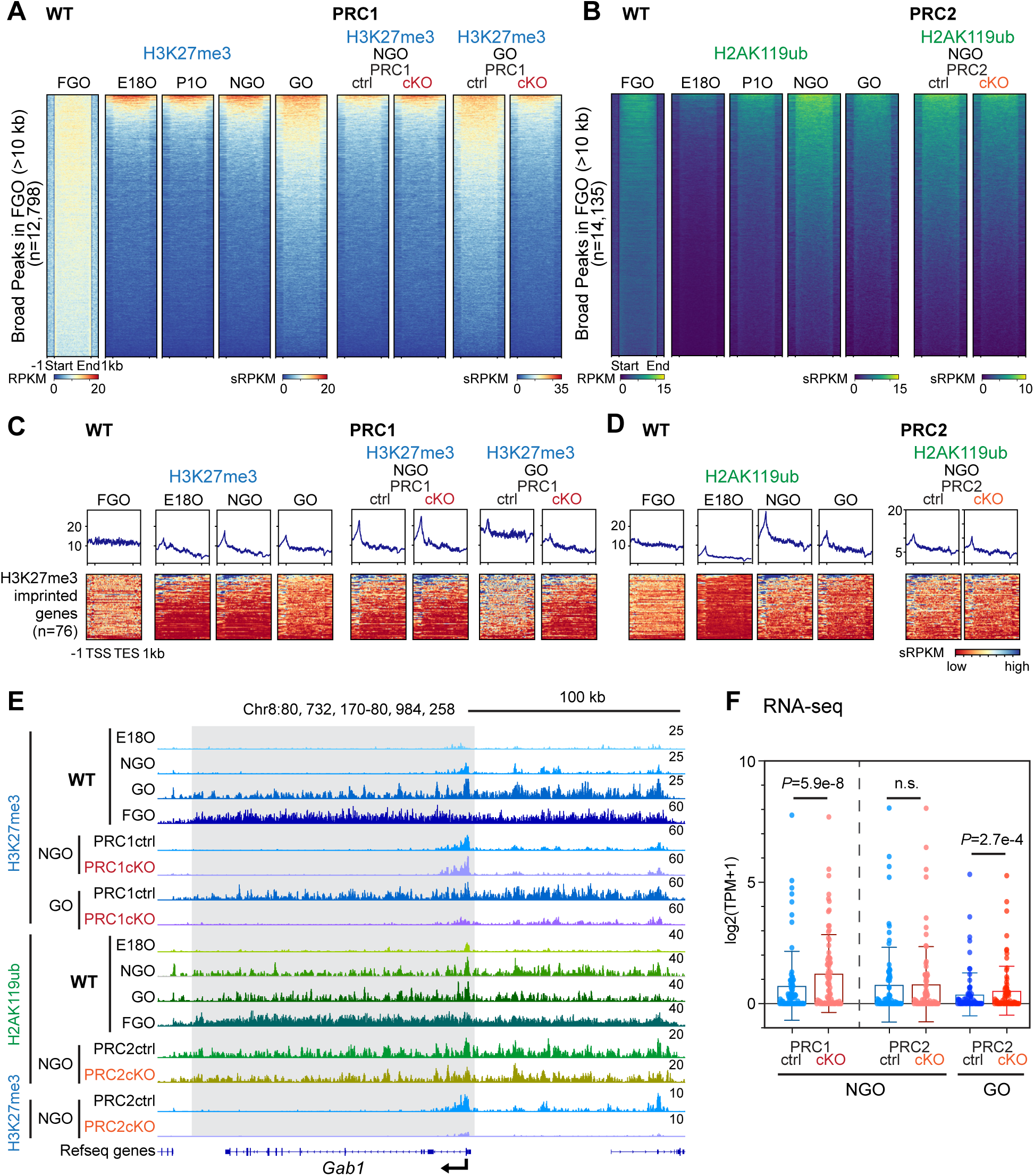
Broad Polycomb domains and H3K27me3-dependant imprinting are primed during perinatal oogenesis. (A) Heatmaps showing H3K27me3 dynamics in WT perinatal oocytes and PRC1ctrl&cKO oocytes on the H3K27me3 broad peaks (peaks length >10kb) identified in FGO. (B) Heatmaps showing H2AK119ub dynamics in WT perinatal oocytes and PRC2ctrl&cKO oocytes on the H2K119ub broad peaks (peaks length >10kb) identified in FGO. (C and D) Heatmaps showing H3K27me3 (C) and H2AK119ub (D) dynamics in WT oocytes, PRC1ctrl& cKO or PRC2ctrl&cKO oocytes on the 76 H3K27me3-imprinted gene regions. (E) Genome browser views showing H2AK119ub and H3K27me3 dynamics at the H3K27me3-imprinted gene *Gab1* locus in WT oocytes and PRC1ctrl&cKO oocytes. (F) Scatter dot plots showing RNA expression levels of the 76 H3K27me3-imprinted genes in PRC1ctrl& cKO or PRC2ctrl&cKO oocytes as indicated. *P* values of pairwise comparisons (two-sided paired Student’s t-test) are given. n.s., not significant. Data are represented as mean ± SD.

Given that maternal H3K27me3 mediates DNA methylation-independent imprinting^21,22,57,58^, we next tested whether H3K27me3-dependent imprinting could be traced back to perinatal oogenesis. At the 76 putative H3K27me3-imprinted gene loci^21^, H3K27me3 coexists with H2AK119ub in FGOs^24,25^, and it has been shown that a variant PRC1 subcomplex is critical for establishing H3K27me3 on these genes^25^. Not known is when H3K27me3 deposition is established. We found that H3K27me3 accumulated in a PRC1-dependent manner at gene bodies in GOs (Figure 6C). H2AK119ub was already present at gene bodies in NGOs independent of PRC2 (Figure 6D). A representative track view at the *Gab1* locus confirms this pattern (Figure 6E). These results suggest a priming mechanism in which PRC1-H2AK119ub sets the stage for the H3K27me3-dependent imprinting in the ovarian reserve. Although the RNA transcripts of these imprinted genes were mostly undetectable or low in oocytes, the loss of PRC1, but not PRC2, resulted in derepression of these genes in NGOs (Figure 6F). Absence of PRC2 leads to a derepression of these genes only in GOs (Figure 6F). Taken together, these results suggest that PRC1 plays a crucial role in regulating H3K27me3 reprogramming during PPT, including the establishment of broad noncanonical domains and H3K27me3 deposition of imprinted genes.

### Polycomb complexes are indispensable for maintenance of ovarian reserve

Finally, we examined the physiological roles of PRC1 and PRC2 *in vivo*. Our previous work demonstrated that PRC1cKO females, driven by *Ddx4*-Cre, exhibited massive oocyte loss during ovarian reserve formation, leading to premature ovarian failure in early adulthood^14^. To further elucidate the role of PRC1 in the long-term maintenance of the ovarian reserve, we generated a new PRC1cKO model using *Gdf9*-iCre^59^, an oocyte-specific Cre recombinase expressed from the primordial follicle stage around P3 (hereafter referred to as PRC1GcKO). However, challenges arose due to the extremely low birth frequency of mutant females from this model, which limited our ability to collect sufficient samples for extensive analysis. We were only able to obtain a total of five PRC1GcKO females for phenotyping. Initial examination at P8, shortly after PRC1 deletion, revealed no obvious differences between the PRC1GcKO ovaries and those of their littermate controls (Figure S10C). However, within two months, a dramatic change was observed; the ovaries of PRC1GcKO mice exhibited significant degeneration, with almost no live oocytes remaining (Figures 7A, B). Notably, clusters of follicle structures devoid of oocytes, indicative of early-stage follicular atresia, were frequently observed in the 2-month-old PRC1GcKO ovaries, a pathology not present in the controls (indicated with asterisks in Figure 7A). These findings provide compelling evidence that PRC1 is critical not only for forming but also for maintaining the ovarian reserve, underscoring its essential role in female germline development.

**Figure 7.**
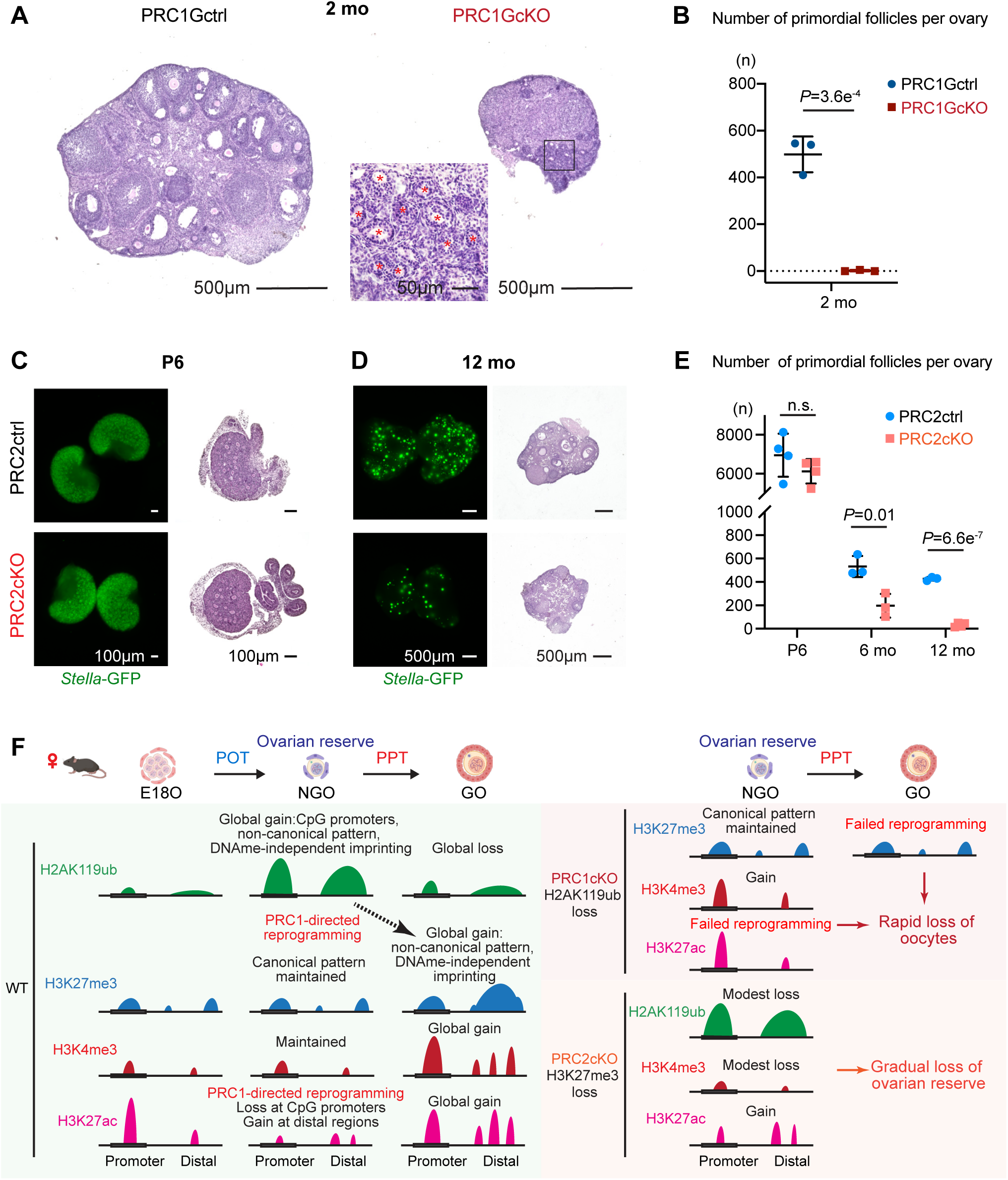
Premature ovarian failure in Polycomb conditional knockout mouse models. (A) Histology of ovarian sections from 2-month-old PRC1Gctrl and PRC1GcKO females, respectively, stained with hematoxylin & eosin. Red asterisks mark atretic follicles. Bars: 500 μm, 50 μm in boxed area. Three mice of each genotype were used for analysis, and representative images are shown. (B) Estimated numbers of primordial follicles per ovary from PRC1Gctrl and PRC1GcKO mice at 2 months. Three mice were analyzed for each genotype. Data are presented as mean values ± SD. Two-tailed unpaired Student’s t-test. (C, D) Ovarian sections of PRC2ctrl and PRC2cKO mice at P6 (C) and 12 months of age (D). The ovaries were directly imaged for *Stella*-GFP (green) or stained with hematoxylin & eosin on sections. Bars: 100 μm for P6 ovaries, 500 μm for 12 mo ovaries. At least three mice were analyzed for each genotype at each time point, and representative images are shown. (E) Estimated numbers of primordial follicles per ovary from PRC2ctrl and PRC2cKO mice at P6, 6 months, and 12 months of age. At least three mice were analyzed for each genotype at each time point. Data are presented as mean values ± SD. n.s., not significant; two-tailed unpaired Student’s t-test. (F) Schematic models of Polycomb-mediated programming and reprogramming in the ovarian reserve development.

We also suspected a role for PRC2-H3K27me3 in ovarian reserve formation based on its relationship with PRC1 and its abundant presence in perinatal oocytes. Surprisingly, ovaries from PRC2cKO mice showed no obvious defects in ovarian reserve formation (Figures 7C, E, and Figure S10D), suggesting that PRC2, unlike PRC1, is not essential for perinatal oogenesis. Given this unexpected finding, we further examined whether PRC2 plays a role in the long-term maintenance of the ovarian reserve. By 6 months, there was a significant loss of primordial follicles in the PRC2cKO ovaries, and by 12 months, the ovarian reserve was nearly depleted, with only a few growing oocytes remaining (Figures 7D, E). These results indicate that, although PRC2 may not be critical for the initial formation of the ovarian reserve or oocyte growth and maturation^57^, it is required for the long-term maintenance of the ovarian reserve.

To investigate potential underlying mechanisms for why PRC2 is required for long-term maintenance of the ovarian reserve, we analyzed the transcriptomes of PRC2cKO oocytes (Figures S10A, B, E, F). We identified 404 genes that were significantly up-regulated and 588 genes that were down-regulated in PRC2cKO NGOs compared to controls (Figure S10B). Although these differences in the transcriptome appear relatively mild, gene ontology analysis offered some insights (Figure S10G). The up-regulated DEGs were enriched with genes involved in meiotic prophase I, such as the meiotic cell cycle/spermatogenesis (e.g., *Sycp1, Hormad1, Majin, M1ap, Spata22, Xlr, Taf7l*), programmed cell death (e.g., *Pmaip1, Phlda2, Foxo1, Id3, Dusp1*), and typical developmental processes. Down-regulated DEGs were highly enriched for genes involved in the regulation of DNA damage response/DNA repair (e.g., *Rad9a, H2ax, Apbb1, Trp53bp1, Ppp4c, Xrcc1, Fanci, Ddx11*), organelle transport, cellular catabolism, etc. Thus, the Gene Ontology analysis suggests that PRC2, like PRC1^14^, plays a role in the exit from the meiotic prophase I program and suppression of ectopic developmental programs and programmed cell death, as well as maintaining the DNA damage/stress response in NGOs.

In summary, we conclude that PRC1 is indispensable for formation of the ovarian reserve, while PRC2 primarily supports its long-term maintenance.

## Discussion

Formation and maintenance of the ovarian reserve is a key determinant of female fertility and reproductive lifespan, but we know little about the molecular mechanisms regulating these processes. Previous studies suggested that epigenetic marks in the ovarian reserve are scarce^12,60^. In contrast, our quantitative profiling identified H2AK119ub as an abundant modification in the ovarian reserve and revealed two key epigenomic transitions that shape female germline development. These transitions highlight the critical role of Polycomb-mediated histone modifications in establishing and maintaining the ovarian reserve and in priming oocytes for growth and oocyte-to-embryo transition (Figure 7F).

The first transition, POT, is marked by establishing a repressive chromatin state essential to maintaining the ovarian reserve. This repressive chromatin state is achieved primarily by H3K27ac reprogramming and rapid accumulation of PRC1-mediated H2AK119ub. PRC1-H2AK119ub is critical for maintaining meiotic arrest and preventing ectopic expression of developmental programs through epigenetic programming^14^, thereby creating a stable epigenomic landscape that preserves genomic integrity as the ovarian reserve is maintained. The second major transition, PPT, is characterized by genome-wide epigenetic reprogramming involving a PRC1-H2AK119ub-directed redistribution of H3K27me3 and an increase in active marks such as H3K4me3 and H3K27ac. This shift from a repressive to a permissive chromatin state enables transcriptional activation required for oocyte growth, accumulation of necessary cellular resources, and eventual meiotic resumption during ovulation. By identifying these epigenetic transitions, this study provides a comprehensive overview of female germline development by filling a major knowledge gap between the epigenetic reprogramming that occurs in PGCs and the epigenetic programming that occurs during oocyte growth.

PRC1 has emerged as a central regulator of both the formation and maintenance of the ovarian reserve. The rapid degeneration observed in PRC1GcKO ovaries underscores the essential role of PRC1 in sustaining the ovarian reserve. In contrast, the PRC2cKO model shows that whereas initial ovarian reserve formation appears unaffected, there is a gradual depletion of the primordial follicle pool over time. This delayed effect highlights a less immediate but nonetheless essential role for PRC2 in maintaining the epigenetic landscape in the ovarian reserve. Given that folliculogenesis remains unaffected after PPT in the PRC2cKO model using *Ddx4*-Cre, similar to the phenotype observed with *Gdf9*-Cre^57^, H3K27me3 reprogramming during PPT primarily prepares oocytes for early embryonic development rather than directly regulating oocyte growth. Our study highlights important differences between the functions of PRC1 and PRC2. PRC1 exerts a broad impact across all developmental stages, whereas PRC2 is more critical for long-term maintenance of the ovarian reserve. This functional divergence between PRC1 and PRC2 is interesting because it differs from what is observed in other cell types, such as embryonic stem cells (ESCs) ^61–63^, PGCs^49,64,65^, and male germline stem cells^46,66,67^, where PRC1 and PRC2 function synergistically to safeguard cell identity and genomic integrity.

Variant PRC1 subcomplexes (vPRC1) are responsible for regulating PRC2-H3K27me3 deposition^68^. Therefore, vPRC1, in particular PRC1.1 and PRC1.6, are promising candidates for the molecular link between PRC1 and PRC2 during PPT. Concordantly, the vPRC1 components *Pcgf1* and *Pcgf6* are significantly up-regulated during PPT (Figure S4). *Pcgf1* and *Pcgf6* are also implicated in H3K27me3-dependent imprinting in mature oocytes^25^. Our study shows that H3K27me3-mediated imprinting and the formation of broad H3K27me3 domains can be traced back to NGOs, and both are regulated by PRC1-H2AK119ub. Thus, our study addresses the origin of H3K27me3-mediated imprinting in the female germline, which is already evident in the perinatal stages at the time of ovarian reserve formation.

We observe that both PRC1 and PRC2 are required to restrict global levels of H3K27ac. Unlike PRC2, which catalyzes H3K27me3 and typically antagonizes H3K27ac^69–74^, PRC1 has a more complex role in modulating H3K27ac. PRC1 not only counteracts H3K27ac formation by inhibiting acetyltransferase recruitment^44^ or by recruiting deacetylases^45^ but also participates in gene activation under certain conditions by promoting H3K27ac^75–78^. During POT, it is notable that H3K27ac occurs at YY1-binding sites (Figure 3I). Because YY1 can function as a PcG protein^79^ by directly interacting with vPRC1 member RYBP (YY1 binding protein)^80^ or YAF2 (YY1-associated factor 2)^81,82^, the results suggest involvement of vPRC1 during POT. Our study determines that PRC1-mediated H3K27ac reprogramming during POT precedes H3K27me3 reprogramming at PPT; thereby, H3K27ac reprogramming is likely a prerequisite for establishing a unique epigenome later during oocyte growth.

In conclusion, our study unveils the complex epigenetic landscape that governs development of the ovarian reserve and establishes Polycomb-mediated programming and reprogramming as critical processes in the ovarian reserve. Our study demonstrates that epigenetic priming in NGOs shapes the epigenetic landscape during oocyte growth and oocyte-to-embryo transition.

## Methods

### Animals

The oocyte-specific reporter *Stella*-GFP transgenic mice were obtained from Dr. M. Azim Surani^83^. Generation of conditionally deficient *Rnf2* mice on a *Ring1^-/-^* background with *Ddx4-Cre* was performed as described previously^14,46,47^. Briefly, PRC1*cKO* mice *Ring1^-/-^; Ring1^F/-^; Ddx4-Cre* were generated from *Ring1^-/-^; Ring1^F/F^* females crossed with *Ring1^-/-^; Ring1^F/+^; Ddx4-Cre* males and PRC1*ctrl* mice used in experiments were *Ring1^-/-^; Rnf2^F/+^; Ddx4-Cre* littermate females. Similarly, PRC1*GcKO* mice *Ring1^-/-^; Ring1^F/F^; Gdf9-Cre* were generated from *Ring1^-/-^; Ring1^F/F^* females crossed with *Ring1^-/-^; Ring1^F/+^; Gdf9-Cre* males and PRC1*Gctrl* mice used in experiments were *Ring1^-/-^; Rnf2^F/F^* littermate females. PRC2*cKO* mice *Eed^F/-^; Ddx4-Cre* were generated from *Eed^F/F^* females crossed with *Eed^F/+^; Ddx4-Cre* males and *PRC2ctrl* mice used in experiments were *Eed^F/+^; Ddx4-Cre* littermate females. *Eed^F/F^* mice were also crossed with the *Stella*-GFP line to get both PRC2*ctrl* and *cKO* mice with the GFP reporter for oocyte collection using FACS. Generation of mutant *Ring1* and *Rnf2* floxed alleles were reported previously^50,84^. The *Eed flox* (B6;129S1-*Eed* ^tm1Sho^/J), *Ddx4-Cre* [FVB-Tg(Ddx4-cre)1Dcas/J], and *Gdf9-Cre* [Tg(Gdf9-iCre)5092Coo/J] mouse lines were purchased from the Jackson Laboratory (022727, 006954 and 011062, respectively). Mice were maintained on a mixed genetic background of FVB and C57BL/6J.

Mice were maintained on a 12:12 light: dark cycle in a temperature and humidity-controlled vivarium (22 ± 2 °C; 40–50% humidity) with free access to food and water in the pathogen-free animal care facility. Mice were used according to the guidelines of the Institutional Animal Care and Use Committee (IACUC protocol no. 21931) at the University of California, Davis.

### Oocyte collection

For WT profiling experiments, the small oocytes (E18.5, P1, P6) from *Stella*-GFP reporter female mice were isolated by FACS, and the early GOs (P7) were manually collected under the stereomicroscope. For PRC1 conditional knockout studies, both NGOs and GOs were manually collected under the stereomicroscope. For PRC2 conditional knockout studies, NGOs were isolated from PRC2ctrl or cKO female pups carrying *Stella*-GFP reporter by FACS, and GOs were manually collected under the stereomicroscope.

Ovaries at different developmental stages were harvested by carefully removing oviducts and ovarian bursa in PBS. Each pair of ovaries were further digested in 200 μl TrypLE™ Express Enzyme (1X) (Gibco, 12604013) supplemented with 0.3% Collagenase Type 1 (Worthington, CLS-1) and 0.01% DNase I (Sigma, D5025) and incubated at 37 for 20-40 min (longer time for bigger postnatal ovaries) with gentle agitation. After incubation, the ovaries were dissociated by gentle pipetting using the Fisherbrand^TM^ Premium Plus MultiFlex Gel-Loading Tips until no visible tissue pieces. Two ml DMEM/F-12 medium (Gibco, 11330107) supplemented with 10% FBS (HyClone, SH30396.03) were then added to the suspension to stop enzyme reaction.

For FACS preparation, the cells were then washed with FACS buffer (PBS containing 2%FBS) twice by centrifugation at 300 × g for 5 min and filtered into a 5 ml FACS tube with a 35 μm nylon mesh cap (Falcon, 352235). Cells were analyzed after removing small and large debris in FSC-A versus SSC-A gating, and doublets in FSC-W versus FSC-H gating. Then, the desired small oocyte population (enriched GFP+ cells with uniform size) was collected based on both GFP signal gating and FSC-A versus SSC-A size backgating.

For manual picking, the cell suspension was filtered through a 100 μm cell strainer and seeded onto a culture dish. The cells were allowed to settle down for 15-30 min at 37 ; 5% CO_2_ before being transferred under the stereomicroscope (Nikon, SMZ1270). Based on morphology and diameter, non-growing oocytes or growing oocytes were specifically picked up, washed, and transferred into the downstream buffer by mouth pipette with a glass capillary.

### Histology and Immunostaining

To prepare ovarian paraffin blocks, ovaries were fixed with 4% paraformaldehyde (PFA) overnight at 4°C. Ovaries were dehydrated and embedded in paraffin. For histological analysis, 5-µm-thick paraffin sections were deparaffinized and stained with hematoxylin and eosin. For immunostaining, ovarian paraffin sections were deparaffinized and autoclaved in target retrieval solution (DAKO) for 10 min at 121°C. Sections were blocked with Blocking One Histo (Nacalai, 06349-64) for 1 h at room temperature and then incubated with primary antibodies (rabbit anti-H3K27me3, 1:100, Cell Signaling Technology, 9733; rabbit anti-DDX4, 1:400, Abcam, ab13840) overnight at 4°C. Localization of the primary antibody was performed by incubation of the sections with the corresponding secondary antibodies (Donkey Anti-Rabbit IgG (H+L) Alexa Fluor 555, Invitrogen, A-31572) at 1:500 dilution for 1h at room temperature. Finally, sections were counterstained with DAPI and mounted using ProLong Gold Antifade Mountant (Invitrogen, P36930). Images were obtained with an ECLIPSE Ti-2 microscope (Nikon) and processed with NIS-Elements (Nikon).

### Quantification of ovarian follicles

Quantification of ovarian follicles was performed as previously described^85^. Briefly, to count the number of follicles, paraffin-embedded ovaries were serially sectioned at 5-μm thickness, and all sections were mounted on slides. Then, these sections were stained with hematoxylin and eosin for morphological analysis. Ovarian follicles at different developmental stages were counted in every fifth section of the collected sections from one ovary, based on the well-accepted standards established by Pederson and Peters^86^. In each section, only those follicles in which the nucleus of the oocyte was clearly visible were scored, and the cumulative follicle counts were multiplied by a correction factor of 5 to represent the estimated number of total follicles in an ovary.

### Quantitative CUT&Tag library generation and sequencing

CUT&Tag libraries of oocytes were prepared as previously described^33,34^ with some modifications (a step-by-step protocol https://www.protocols.io/view/bench-top-cut-amp-tag-kqdg34qdpl25/v3) using CUTANA™ pAG-Tn5 (Epicypher, 15-1017). To perform quantitative spike-in CUT&Tag, Drosophila S2 cells were added to mouse oocytes at a fixed ratio (e.g., 2000 S2 cells to 1000 mouse oocytes) at the beginning of each reaction. The antibodies used were rabbit anti-H2AK119ub (1/100; Cell Signaling Technology; 8240), rabbit anti-H3K27me3 (1/50; Cell Signaling Technology; 9733), rabbit anti-H3K4me3 (1/50; Active Motif; 39159) and rabbit anti-H3K27ac (1/50; Cell Signaling Technology; 8173). CUT&Tag libraries were sequenced on the HiSeq 4000 system (Illumina) with 150-bp paired-end reads.

### RNA-seq library generation and sequencing

RNA-seq libraries of PRC2ctrl and cKO oocytes were prepared as follows. ∼750 non-growing oocytes or ∼150 growing oocytes from P6-7 ovaries were collected as one replicate, and two independent biological replicates were used for RNA-seq library generation. Total RNA was extracted using the RNeasy Plus Micro Kit (QIAGEN, Cat # 74034) according to the manufacturer’s instructions. Library preparation was performed with NEBNext® Single Cell/Low Input RNA Library Prep Kit for Illumina® (NEB, E6420S) according to the manufacturer’s instructions. Prepared RNA-seq libraries were sequenced on the HiSeq 4000 system (Illumina) with paired-ended 150-bp reads.

### RNA-seq data processing

Raw RNA-seq reads after trimming by Trim-galore (https://github.com/FelixKrueger/TrimGalore) (version 0.6.10) with the parameter ‘--paired’ were aligned to the mouse (GRCm38/mm10) genome using HISAT2 ^87^ (version 2.2.1) with default parameters. All unmapped reads, non-uniquely mapped reads, and reads with low mapping quality (MAPQ < 30) were filtered out by samtools ^88^ (version 1.16.1) with the parameter ‘-q 30’ before being subjected to downstream analyses.

To identify differentially expressed genes in RNA-seq, raw read counts for each gene were generated using the featureCounts^89^ part of the Subread package (version 2.0.3) based on mouse gene annotations (gencode.vM25.annotation.gtf, GRCm38/mm10). DESeq2^90^ (version 1.44.0) was used for differential gene expression analyses with cutoffs FoldChange > 1.5 and P_adj_ values < 0.05. In DESeq2, P values attained by the Wald test were corrected for multiple testing using the Benjamini and Hochberg method by default. P_adj_ values were used to determine significantly dysregulated genes. GO analyses were performed using the online functional annotation clustering tool Metascape ^91^ (http://metascape.org). The TPM values of each gene were generated using RSEM ^92^(version 1.3.3) for comparative expression analyses and computing the Pearson correlation coefficient between biological replicates. RNA expression heatmaps were plotted using Morpheus (https://software.broadinstitute.org/morpheus/).

Raw RNA-seq datasets of WT oocytes downloaded from GSE128305 (ref.^30^) were processed with the same pipeline with modifications adapting single-end reads. For differential gene expression analyses, stringent cutoffs FoldChange > 2 and P_adj_ values < 0.05 were applied.

### CUT&Tag data processing

#### Upstream processing

We basically followed the online tutorial posted by the Henikoff Lab (https://protocols.io/view/cut-amp-tag-data-processing-and-analysis-tutorial-bjk2kkye.html) with some modifications. Briefly, after trimming by Trim-galore (https://github.com/FelixKrueger/TrimGalore) (version 0.6.10), raw paired-end reads were aligned with the mouse genome (GRCm38/mm10) using Bowtie2 ^93^ (version 2.4.5) with options: --end-to-end --very-sensitive --no-mixed --no-discordant --phred33 -I 10 -X 700. *D. melanogaster* DNA delivered by *Drosophila* S2 cells was used as spike-in DNA for CUT&Tag. For mapping *D. melanogaster* spike-in fragments, we also use the --no-overlap --no-dovetail options to avoid cross-mapping. PCR duplicates from all mapped bam files were removed using the ‘MarkDuplicates’ command in Picard tools (version 3.0.0) (https://broadinstitute.github.io/picard/). To compare replicates, Pearson correlation coefficients were calculated and plotted by ‘multiBamSummary bins’ and ‘plot correlation’ commands from deepTools ^94^ (version 3.5.5). Biological replicates were pooled using the ‘merge’ command in samtools for visualization and other analyses after confirming reproducibility (Pearson correlation coefficient > 0.85). Spike-in normalization was implemented using the exogenous scaling factor computed from the dm6 mapping files (scale factors = 10,000 / aligned spike-in reads).

#### Downstream analysis and visualization

Spike-in normalized genome coverage tracks with 1 bp resolution in BigWig format were generated using ‘bamCoverage’ from deepTools with the parameters ‘--binSize 1 --extendReads --samFlagInclude 64 --normalizeUsing RPKM -- scaleFactor $scale_factor’. Track views were visualized and exported from Integrative Genomics Viewer ^95^ (version 2.14.1). Average scores (sRPKM: scaled RPKM) per region [10kb bins or custom intervals, e.g., peaks, promoters (TSS±1kb)] were calculated from scaled BigWig files using the ‘multiBigwigSummary bins’ or ‘multiBigwigSummary BED-file’ functions from deepTools with adding parameter ‘--outRawCounts’. Blacklisted regions were excluded from analyses by applying the parameter ‘--blackListFileName mm10-blacklist.v2.bed’ while using deepTools. This output matrix from deepTools is used for the following comparative analyses. Hierarchical clustering analysis was conducted in Morpheus using sRPKM values of each 10kb bin across the entire genome as input. Sum sRPKM values of all bins from each replicate track were used to calculate relative global levels of histone modifications. Following a previous study^96^, average sRPKM values of 10kb bins or custom intervals are transformed to integers and used as input for differential enrichment analyses, performed by DESeq2 with fixed size factors since BigWig files were already spike-in normalized. Differential enriched regions are identified with cutoffs log2FoldChange > 1 and P_adj_ < 0.05. Plots were created using the R package ggplot2 ^97^ (v3.5.1).

Bivalent promoters were characterized based on the enrichment level (sRPKM values) of H3K4me3 and H3K27me3 on promoter regions (TSS±1kb) calculated by deepTools. All the promoters (H3K4me3 sRPKM>20 & H3K27me3 sRPKM>2) were defined as bivalent promoters. The cutoff values were confirmed by manual visualization of the track views.

Peak calling for H2AK119ub and H3K27me3 was performed using the Sparse Enrichment Analysis for CUT&RUN (SEACR) ^98^ (https://seacr.fredhutch.org/) by selecting the top 5% enriched regions in stringent mode with parameters’-c 0.05 -n norm -m stringent’. MACS2^99^ was used for peak calling of H3K4me3 and H3K27ac with the parameters’-p 1e-5 -- broad -g mm’. HOMER ^100^ (version 4.11) was used for peak annotation and finding motifs. Peaks comparison was done by the ‘intersect’ command of bedtools ^101^ (version 2.31.0). For differential enrichment analyses on peaks, peaks of each genotype or stage were merged by BEDtools and then used as the reference genomic intervals for counting. ChIP-x Enrichment Analysis (ChEA) was performed using the Enricher website (https://maayanlab.cloud/Enrichr/) ^102,103^. DeepTools was used to draw tag density plots and heatmaps for enrichments.

### Chromatin states analysis using ChromHMM

The ChromHMM^35^ software (version 1.25) was used to classify genomic regions based on the dynamics of all four histone modifications in perinatal oogenesis. Specifically, signal files in bed format (transformed from above scaled Bigwig files) for each histone modification from all stages were first binarized using the ‘BinarizeBed’ command with a bin size of 5 kb (option ‘-b 5000’). The binarized signal files were then used as input to characterize chromatin states conducted using the ‘LearnModel’ command with 5-kb bins (options’-b 5000’). The number of states to include in the mode was finally set to 12 for best visualization after comparing outputs for different numbers.

### Identification of CGI genes and non-CGI genes

CpG islands (CGIs) were defined as published^104,105^, and their coordinates of mouse genome GRCm38/mm10 assembly (n=17,017) were downloaded from UCSC genome browser (https://genome.ucsc.edu/cgi-bin/hgTables?db=mm10&hgta_group=regulation&hgta_track=cpgIslandExt&hgta_table=cpgIslandExt&hgta_doSchema=describe+table+schema). Genes were classified as CGI-associated if a first base pair of any transcript TSSs ±100-bp overlapped a CGI. In this way, the total 53,795 genes from Gencode vM25 mouse gene annotations (after removing Chromosome Y and M genes) were divided into CGI genes (n=14,040) and non-CGI genes (n=39,755).

### CUT&RUN data reanalysis

Raw CUT&RUN data of H2AK119ub and H3K27me3 in GV FGOs were downloaded from Gene Expression Omnibus (GEO) under accession no. GSE153529 (ref. ^24^). Raw reads files were processed using the same pipeline as CUT&Tag datasets for alignment and duplicate removal. Deduped bam files were converted to bigwig files using deeptools by normalizing with RPKM or RPGC and then used for plotting. Peak calling for H2AK119ub and H3K27me3 was performed using the SEACR by selecting the top 5% enriched regions in stringent mode with parameters’-c 0.05 -n norm -m stringent’. Peaks (width>10kb) were selected as broad domains and used for plotting.

### WGBS data reanalysis

Raw WGBS data of GV FGOs were downloaded from DDBJ: DRA000570 (ref. ^8^). Low-quality bases at the 3’ end and residual adapter sequences were removed from the raw sequencing reads. The processed reads were then aligned to the mouse genome (mm10) using Bismark with the PBAT option. Bisulfite conversion rates were estimated from reads uniquely aligned to the lambda phage genome. Counts from the two cytosines in a CpG dinucleotide and its reverse complement were combined, and only CpG sites with coverage between 5× and 200× were included in the analysis of CpG methylation levels.

### Statistics

Statistical methods and P values for each plot are listed in the figure legends and/or in the Methods. Statistical significance for pairwise comparisons was determined using two-tailed unpaired t-tests. Next-generation sequencing data (RNA-seq, CUT&Tag) were based on two independent replicates. No statistical methods were used to initially determine sample size in these experiments. Experiments were not randomized, and investigators were not blinded to allocation during experiments and outcome assessments.

## Supporting information

Supplemental Figrues

## Resource availability

RNA-seq datasets of WT oocytes were downloaded from GSE128305 (ref.^30^). The H3K27me3 and H2AK119ub CUT&RUN datasets of GV full-grown oocytes were from GSE153529 (ref. ^24^). The DNA methylation WGBS data of GV full-grown oocytes were from DDBJ: DRA000570 (ref. ^8^).

## Acknowledgments

We thank M. Azim Surani for sharing *Stella*-GFP transgenic mice. We thank Yao Cai and Prof. Joanna Chiu from UC Davis for sharing Drosophila S2 cells. Funding sources: Open Collective Foundation Repro Grants (M.H), National Institutes of Health grants R35GM141085 (S.H.N) and R21HD110146 (S.H.N and R.M.S).

## Author contributions

M.H. and S.H.N. designed the study. M.H., Y.H.Y., H.W. performed experiments. M.H., Y.M. and S.H.N. designed and interpreted the computational analyses. M.H., N.H., R.M.S., and S.H.N. interpreted the results and wrote the manuscript with critical feedback from all other authors. R.M.S. and S.H.N. supervised the project.

