## Supplemental Figrues for "An Epigenomic Roadmap Primes Non-Growing Oocytes for Maturation and Early Embryogenesis"

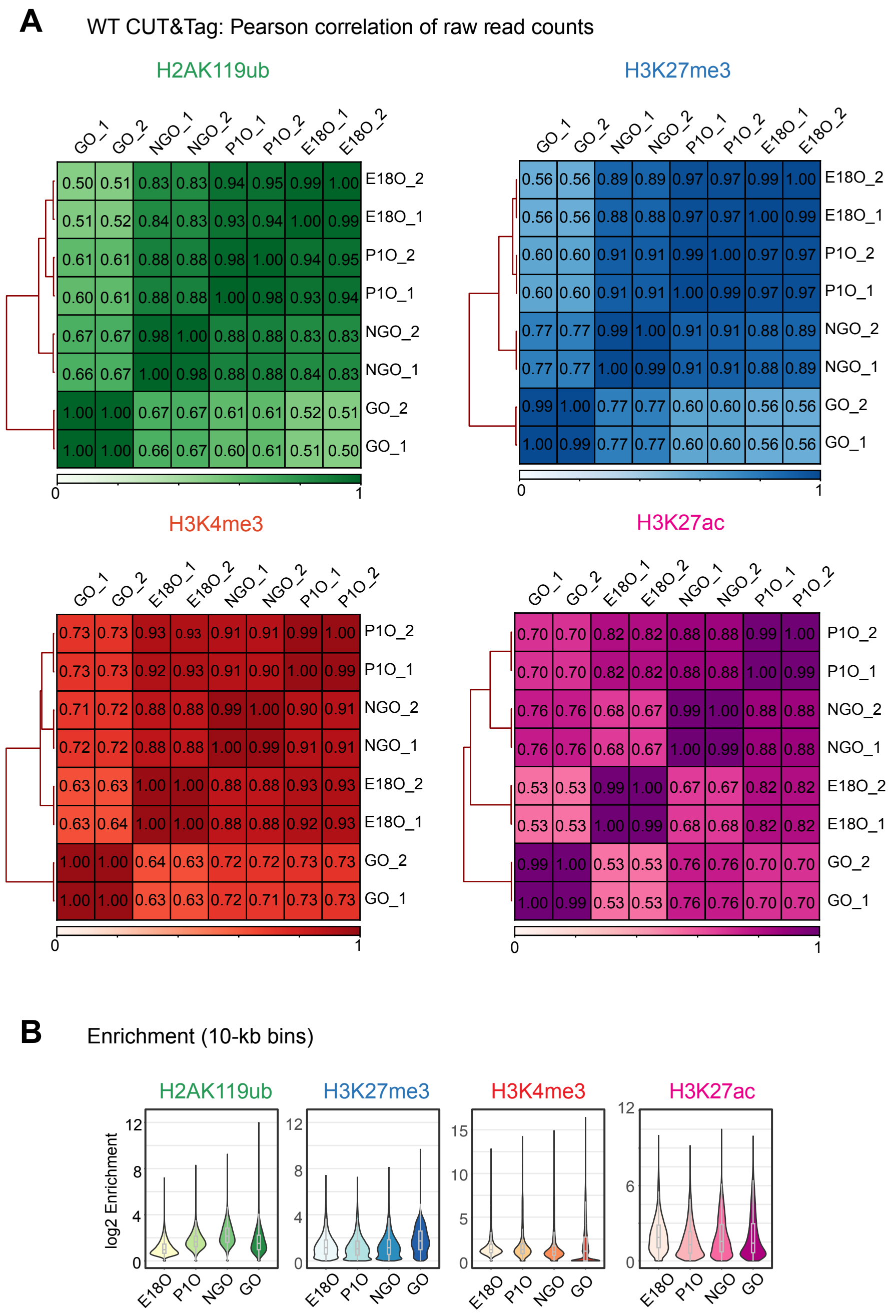


**Figure S1. CUT&Tag datasets of WT oocytes.**
(A) Heatmaps with hierarchical clustering showing the Pearson correlation of the raw read counts among each biological replicate in WT CUT&Tag data.
(B) Violin plots with included boxplots showing average enrichment of H2AK119ub, H3K27me3, H3K4me3, and H3K27ac in 10-kb bins during perinatal oogenesis. Boxes show the 25th and 75th percentile with the median, and whiskers indicate 1.5 times the interquartile range.

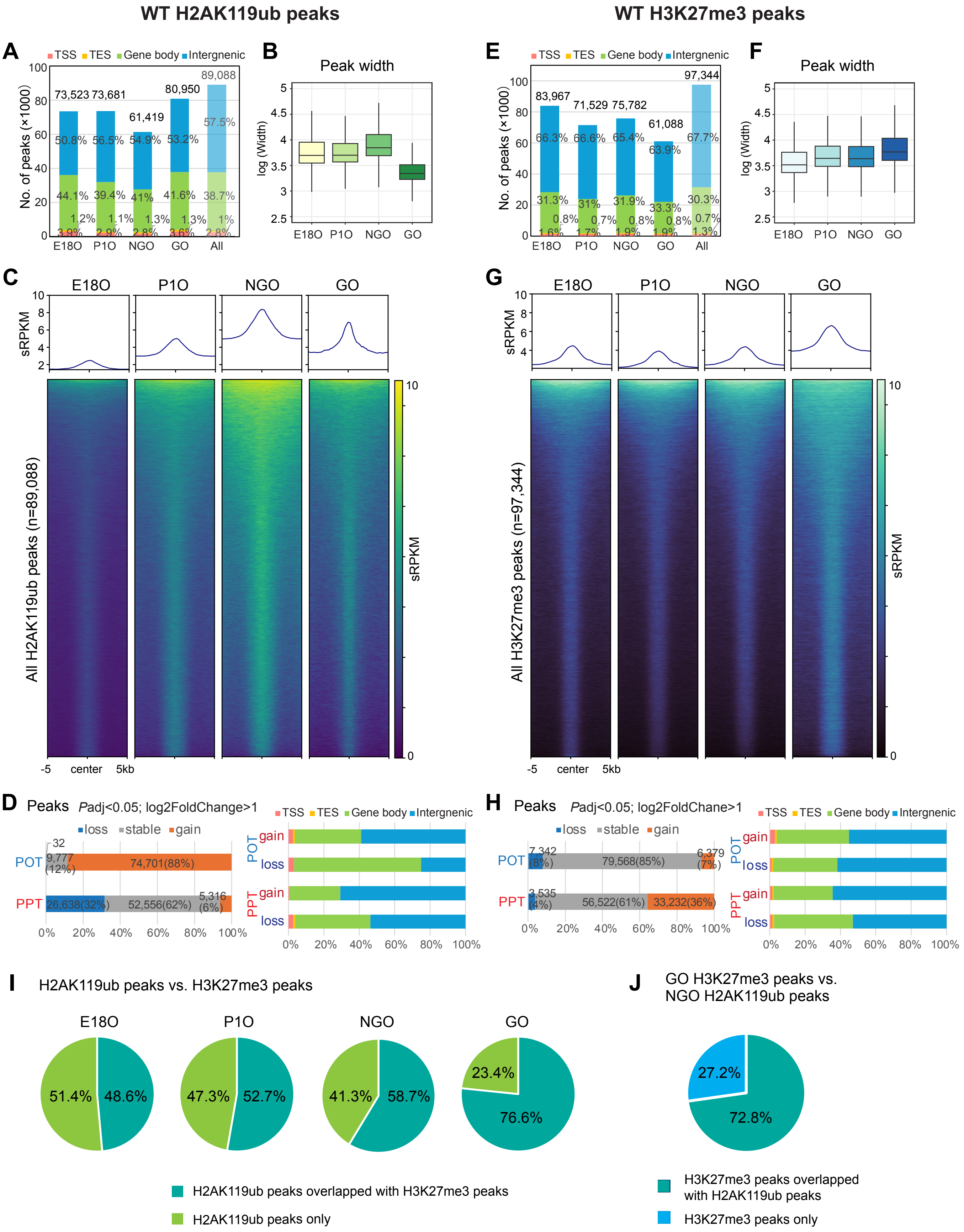
WT H2AK119ub peaks
WT H3K27me3 peaks
A
TSS TES Gene body Intergnenic
No. of peaks (×1000)
73,523 73,681 61,419 80,950 89,088
E18O P1O NGO GO All
B
Peak width
log (Width)
E18O P1O NGO GO
C
E18O P1O NGO GO
sRPKM
All H2AK119ub peaks (n=89,088)
-5 center 5kb
D
Peaks Padj<0.05; log2FoldChange>1
loss stable gain
POT
PPT
E
83,967 71,529 75,782 61,088 97,344
F
Peak width
G
All H3K27me3 peaks (n=97,344)
H
Peaks Padj<0.05; log2FoldChane>1
I
H2AK119ub peaks vs. H3K27me3 peaks
E18O P1O NGO GO
51.4% 48.6%
47.3% 52.7%
41.3% 58.7%
23.4% 76.6%
H2AK119ub peaks overlapped with H3K27me3 peaks
H2AK119ub peaks only
J
GO H3K27me3 peaks vs. NGO H2AK119ub peaks
27.2% 72.8%
H3K27me3 peaks overlapped with H2AK119ub peaks
H3K27me3 peaks only

**Figure S2. Dynamics of Polycomb-mediated repressive marks during perinatal oogenesis.**
(A, E) Bar charts showing H2AK119ub and H3K27me3 peak number and genomic distribution in WT perinatal oocytes.
(B, F) Boxplots showing H2AK119ub and H3K27me3 peak width in WT perinatal oocytes. Boxes show the 25th and 75th percentile with the median, and whiskers indicate 1.5 times the interquartile range.
(C, G) Heatmaps and average tag density plots showing all H2AK119ub and H3K27me3 peaks in WT perinatal oocytes.
(D, H) Bar charts showing percentages of differentially enriched peaks for H2AK119ub and H3K27me3 during POT and PPT and their corresponding genomic distribution.
(I) Pie charts showing percentages of H2AK119ub peaks overlapping H3K27me3 peaks in each stage of perinatal oogenesis.
(J) Pie chart showing the percentage of H3K27me3 peaks in WT GO overlapping H2AK119ub peaks in WT NGO.

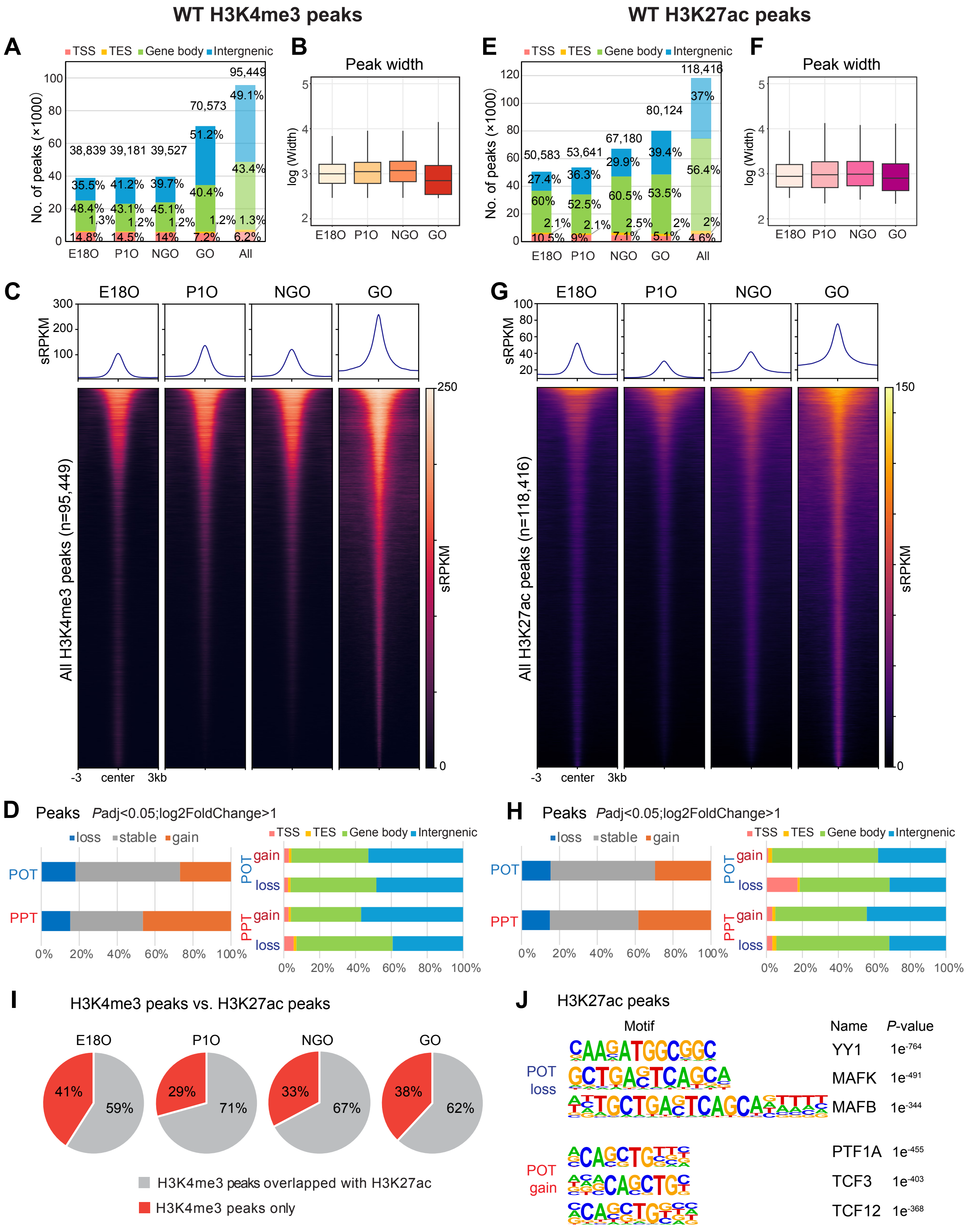

WT H3K4me3 peaks
WT H3K27ac peaks
A
TSS TES Gene body Intergenic
No. of peaks (×1000)
38,839 39,181 39,527 70,573 95,449
E18O P1O NGO GO All
B
Peak width
log (Width)
E18O P1O NGO GO
C
E18O P1O NGO GO
sRPKM
All H3K4me3 peaks (n=95,449)
-3 center 3kb
D
Peaks Padj<0.05;log2FoldChange>1
loss stable gain
POT
PPT
gain
loss
E
No. of peaks (×1000)
50,583 53,641 67,180 80,124 118,416
F
Peak width
G
All H3K27ac peaks (n=118,416)
H
Peaks Padj<0.05;log2FoldChange>1
I
H3K4me3 peaks vs. H3K27ac peaks
E18O P1O NGO GO
41% 59% 29% 71% 33% 67% 38% 62%
H3K4me3 peaks overlapped with H3K27ac
H3K4me3 peaks only
J
H3K27ac peaks
Motif
Name
P-value
POT loss
YY1 1e-764
MAFK 1e-491
MAFB 1e-344
POT gain
PTF1A 1e-455
TCF3 1e-403
TCF12 1e-368

**Figure S3. Dynamics of active marks H3K4me3 and H3K27ac during perinatal oogenesis.**

(A, E) Bar charts showing H3K4me3 and H3K27ac peak number and genomic distribution in WT perinatal oocytes.

(B, F) Boxplots showing H3K4me3 and H3K27ac peak width in WT perinatal oocytes. Boxes show the 25th and 75th percentile with the median, and whiskers indicate 1.5 times the interquartile range.

(C, G) Heatmaps and average tag density plots showing all H3K4me3 and H3K27ac peaks in WT perinatal oocytes.

(D. H) Bar charts showing percentages of differentially enriched peaks for H3K4me3 and H3K27ac during POT and PPT and their corresponding genomic distribution.

(I) Pie chart showing percentages of H3K4me3 peaks overlapping H3K27ac peaks in each stage of perinatal oogenesis.

(J) Motif analysis of differentially enriched H3K27ac peaks during POT (*P* value, hypergeometric test with Bonferroni correction, one-sided, from HOMER).

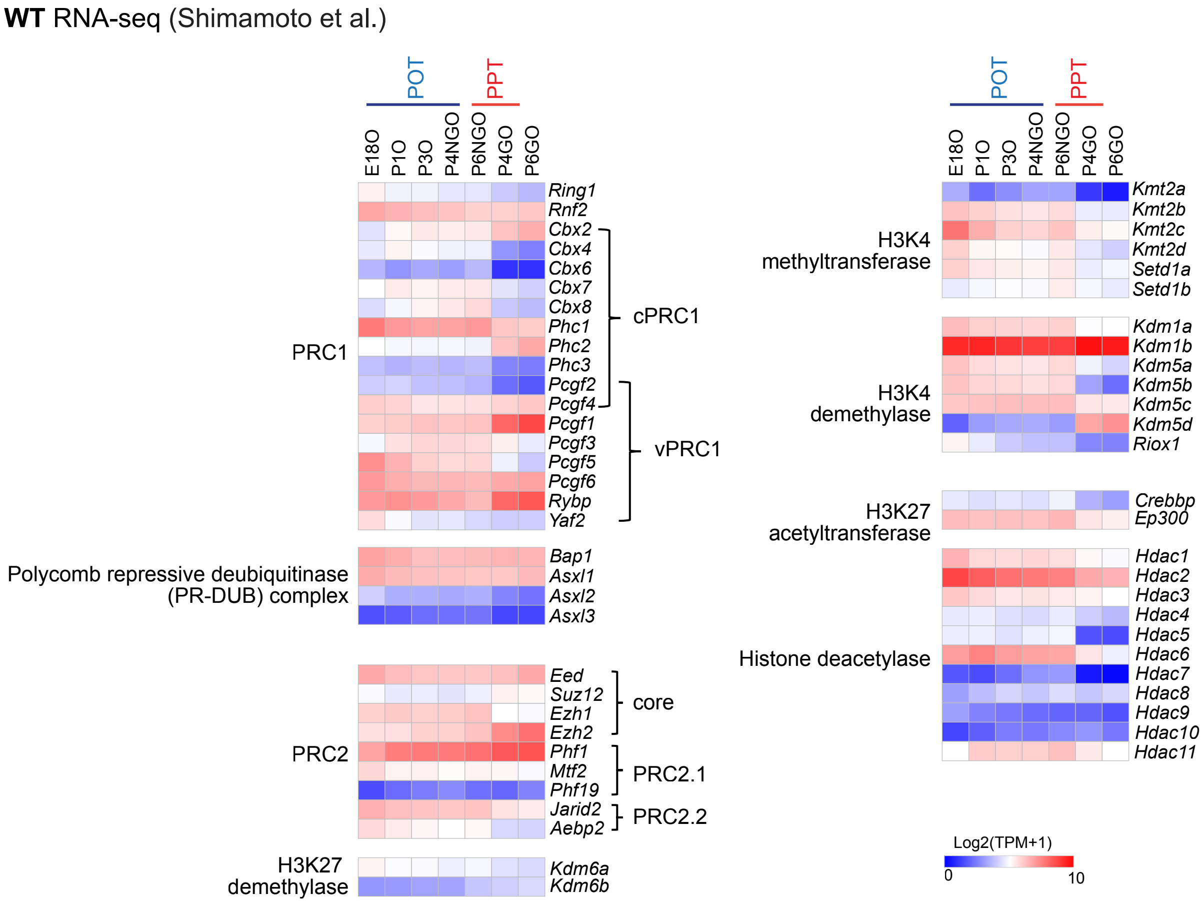


**Figure S4. Expression of chromatin-modifying enzymes related to the four histone modifications.** Heatmap showing gene expression of the key components comprising enzymes related to the four key histone modifications profiled in this study during perinatal oogenesis. In wild-type, E18O represents oocytes in MPI; P1O and P3O represent oocytes transitioning to dictyate arrest;P4 and P6 small oocytes represent NGOs residing in primordial follicles; and P4 and P6 large oocytes represent GOs in primary follicles after initiation of oocyte growth. RNA-seq data of WT oocytes were downloaded from GSE128305 (ref.[31]) and processed. TPM values of genes were used to plot the heatmaps.

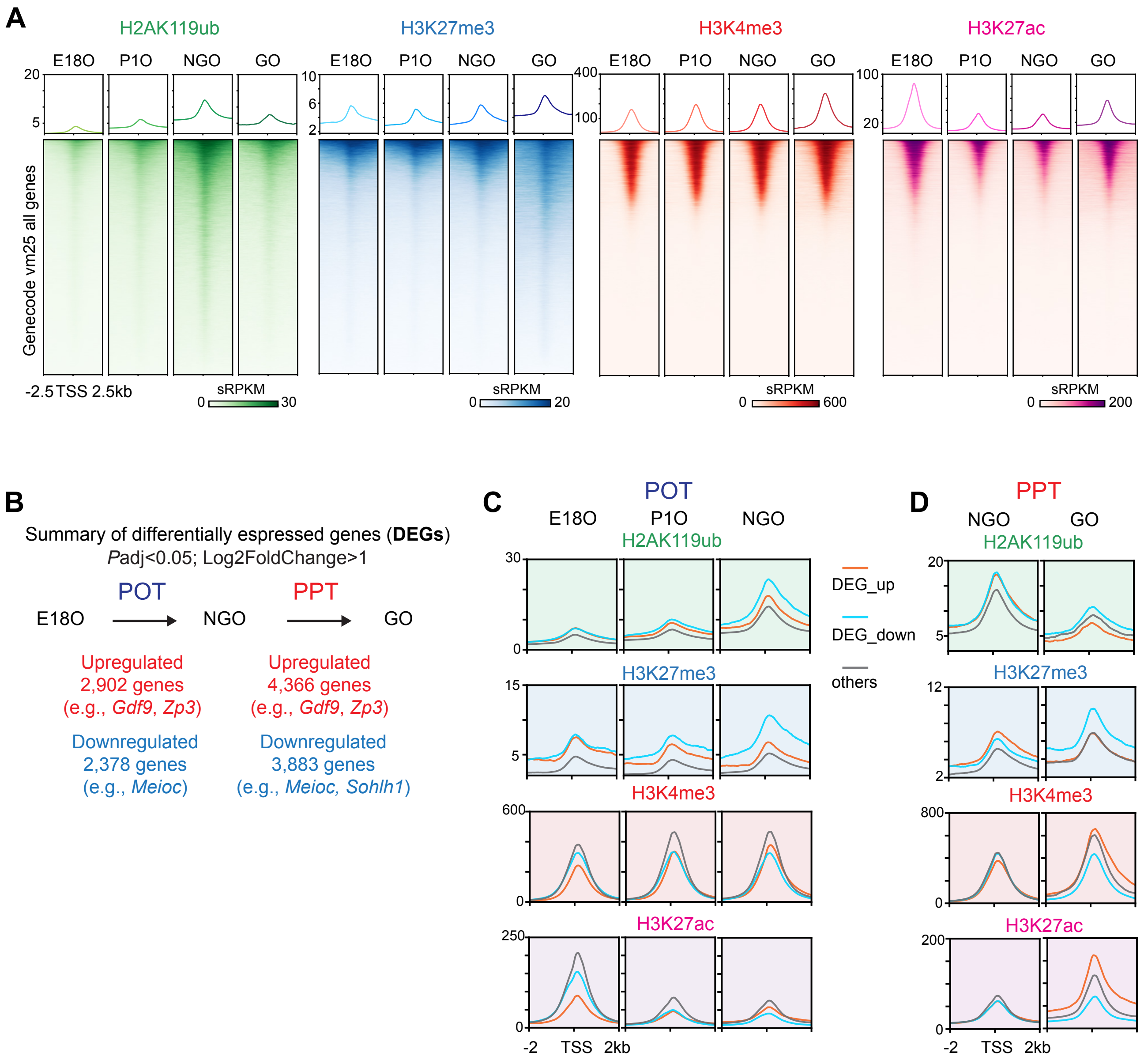


**Figure S5. Histone modifications at gene promoters and differentially expressed genes.**
(A) Average tag density plots and heatmaps showing H2AK119ub, H3K27me3, H3K4me3, and H3K27ac dynamics at promoter regions (TSS ± 2.5 kb) in perinatal oocytes. The color keys represent signal intensity, and the numbers represent spike-in scaled RPKM (sRPKM) values.
(B) Summary of differentially expressed genes (DEGs) during POT and PPT identified in WT oocytes.
(C, D) Average tag density plots showing H2AK119ub, H3K27me3, H3K4me3, and H3K27ac enrichment at promoter regions (TSS ± 2 kb) of DEGs during POT and PPT in WT mice.

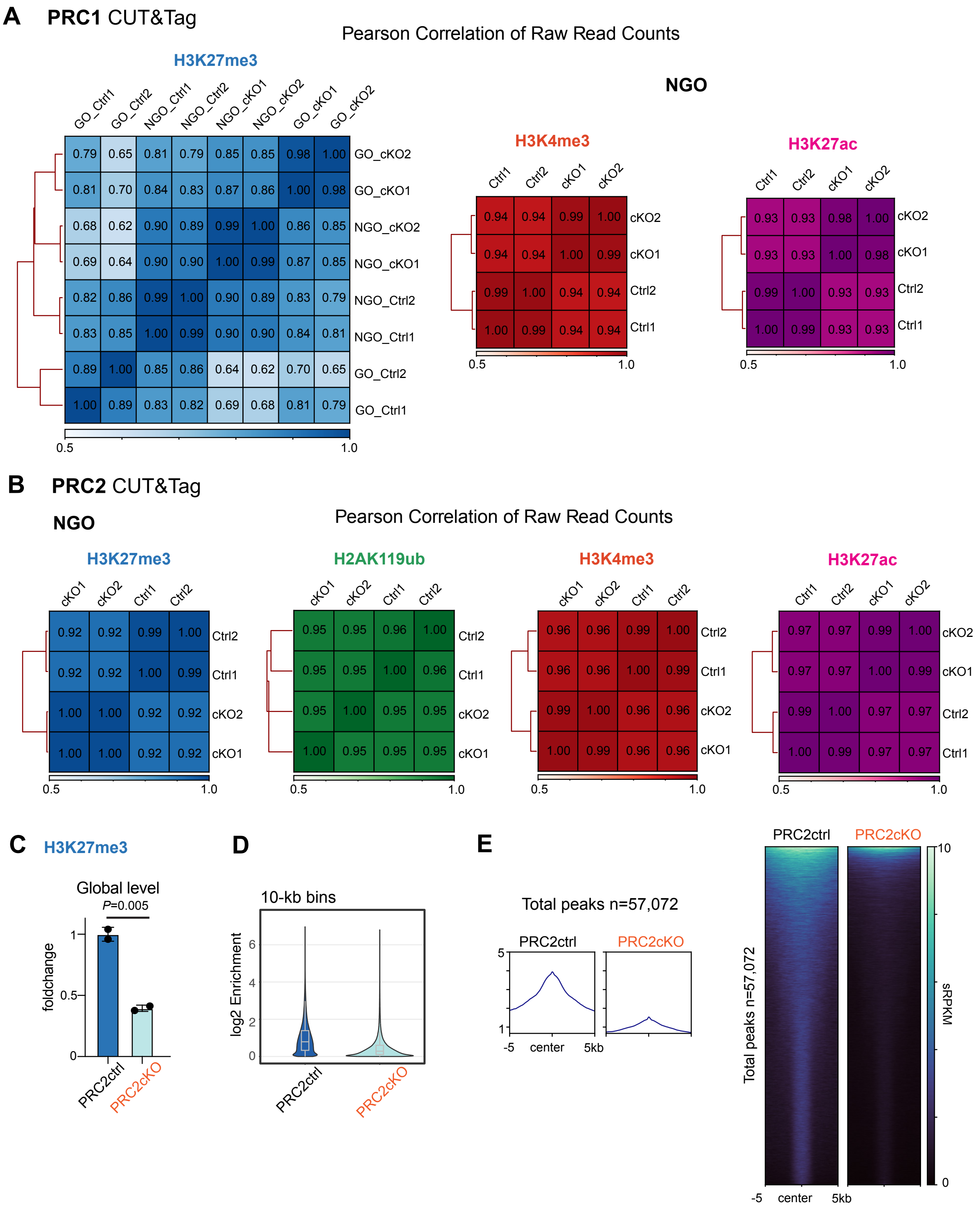

A
PRC1 CUT&Tag
Pearson Correlation of Raw Read Counts
H3K27me3
NGO
H3K4me3
H3K27ac
B
PRC2 CUT&Tag
NGO
Pearson Correlation of Raw Read Counts
H3K27me3
H2AK119ub
H3K4me3
H3K27ac
C
H3K27me3
Global level
P=0.005
foldchange
PRC2ctrl
PRC2cKO
D
10-kb bins
log2 Enrichment
PRC2ctrl
PRC2cKO
E
Total peaks n=57,072
PRC2ctrl
PRC2cKO
-5 center 5kb
Total peaks n=57,072
sRPKM

**Figure S6. CUT&Tag datasets of the Polycomb conditional knockout mouse models.**
(A, B) Heatmaps with hierarchical clustering showing the Pearson correlation of the raw read counts among each biological replicate for each histone modification in CUT&Tag datasets of PRC1ctrl&cKO and PRC2ctrl&cKO, respectively.
(C) Bar chart showing the foldchange of H3K27me3 global level in PRC2ctrl&cKO NGOs (n = 2 biological replicates, indicated by dots). *P* values of pairwise comparisons (two-sided unpaired Student's t-test) are given. Data are represented as mean ± SD.
(D) Violin plots with included boxplots showing average enrichment of H3K27me3 in 10-kb bins in PRC2ctrl&cKO NGOs. Boxes show the 25th and 75th percentile with the median, and whiskers indicate 1.5 times the interquartile range.
(E) Average tag density plots and heatmaps showing all H3K27me3 peaks in PRC2ctrl and PRC2cKO NGOs.

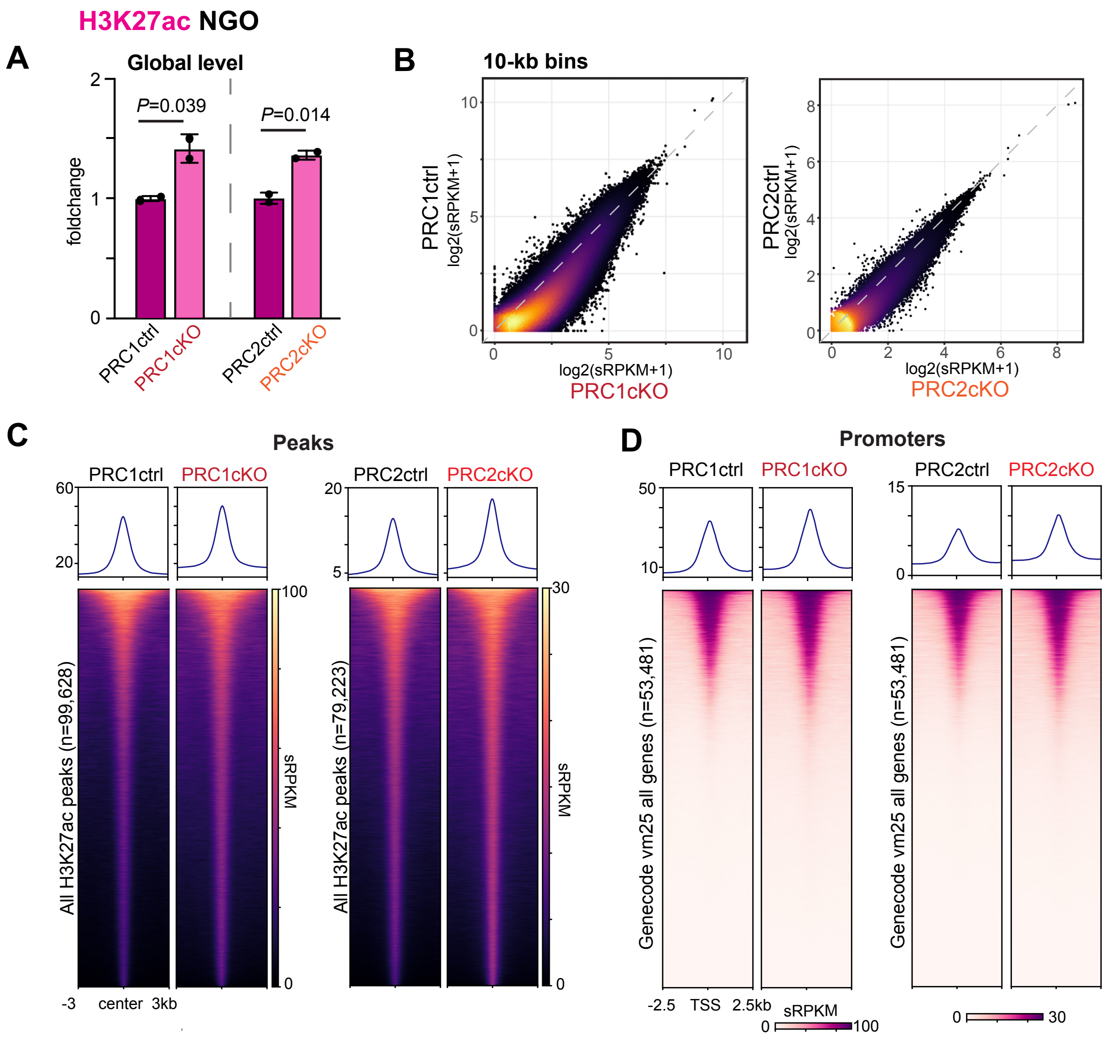


**Figure S7. H3K27ac changes in the Polycomb conditional knockout mouse models.**

(A) Bar chart showing the foldchange of H3K27ac global level in PRC1ctrl&cKO and PRC2ctrl&cKO NGOs (n = 2 biological replicates, indicated by dots). *P* values of pairwise comparisons (two-sided unpaired Student's t-test) are given. Data are represented as mean ± SD.

(B) Density scatter plots showing genome-wide H3K27ac enrichment by 10-kb bins, comparing PRC1ctrl&cKO, PRC2ctrl&cKO, respectively.

(C) Average tag density plots and heatmaps showing all H3K27ac peaks in PRC1ctrl&cKO and PRC2ctrl&cKO NGOs.

(D) Average tag density plots and heatmaps showing H3K27ac enrichment on all promoters in PRC1ctrl&cKO and PRC2ctrl&cKO NGOs.

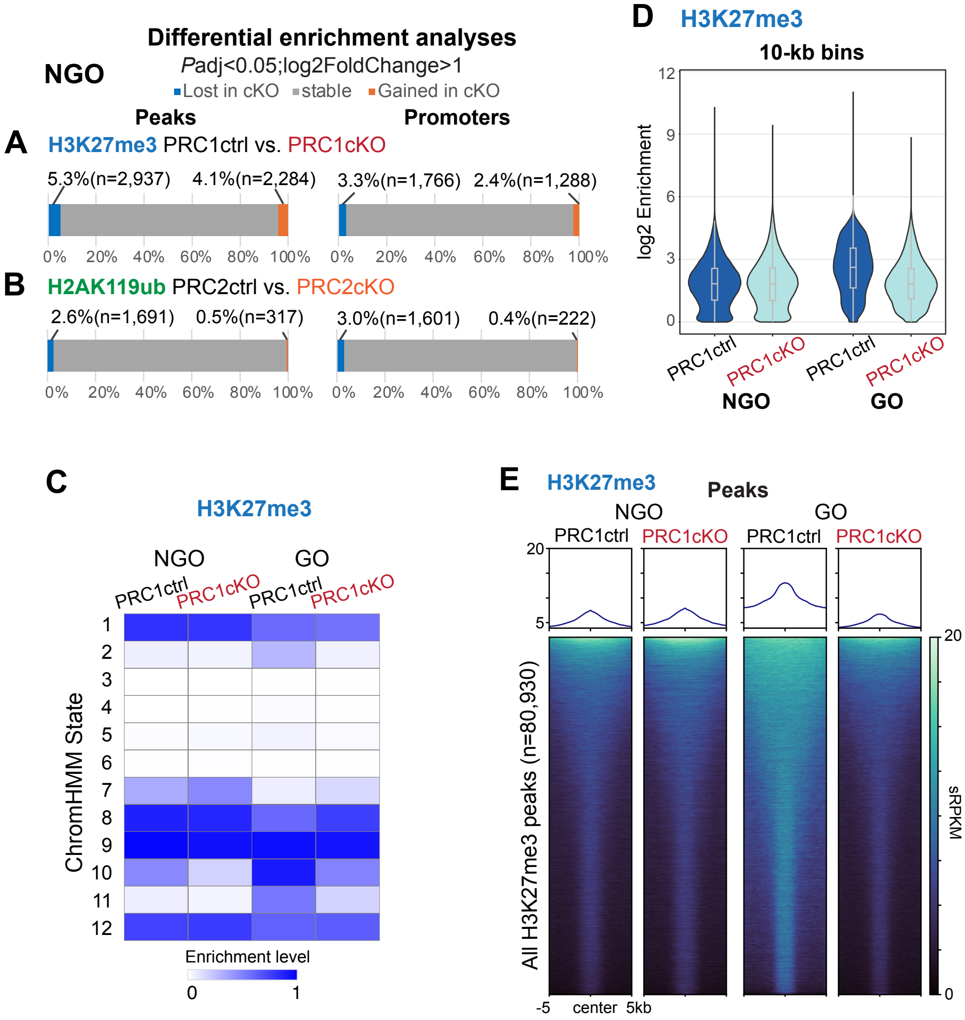


**Figure S8. H2AK119ub and H3K27me3 changes in the Polycomb conditional knockout mouse models.**

(A, B) Bar charts showing percentages of differentially enriched peaks and promoters for H3K27me3 in PRC1ctrl&cKO NGO and H2AK119ub in PRC2ctrl&cKO NGO, respectively.

(C) Heatmap illustrating H3K27me3 levels in PRC1ctrl and PRC1cKO oocytes by chromatin states identified in WT perinatal oocytes using ChromHMM analysis (Figure 1F).

(D) Violin plots with included boxplots showing average enrichment of H3K27me3 in 10-kb bins in PRC1ctrl&cKO oocytes. Boxes show the 25th and 75th percentile with the median, and whiskers indicate 1.5 times the interquartile range.

(E) Average tag density plots and heatmaps showing all H3K27me3 peaks in PRC1ctrl&cKO oocytes.

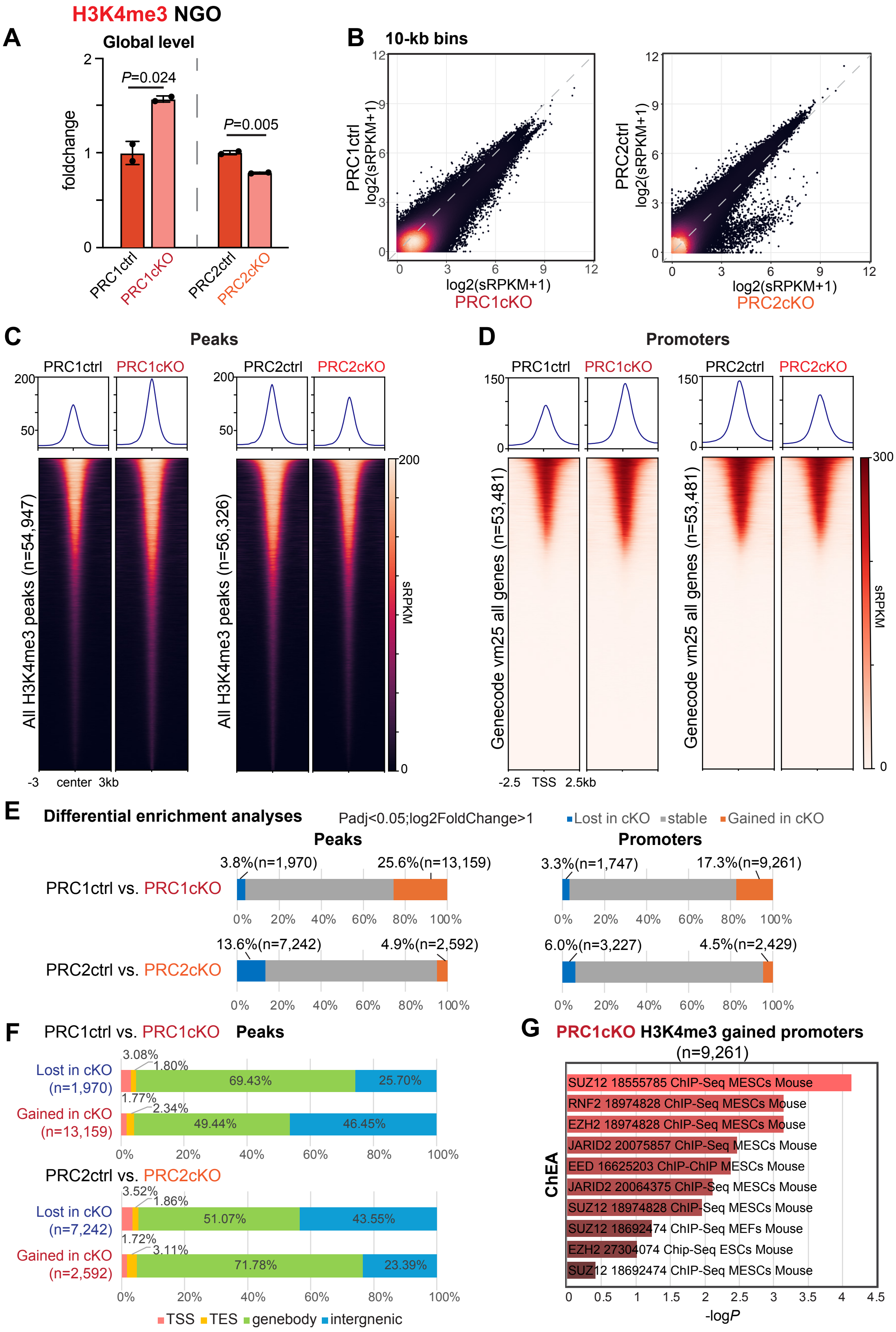

H3K4me3 NGO
A
Global level
foldchange
P=0.024
P=0.005
PRC1ctrl
PRC1cKO
PRC2ctrl
PRC2cKO
B
10-kb bins
PRC1ctrl log2(sRPKM+1)
log2(sRPKM+1)
PRC1cKO
PRC2ctrl log2(sRPKM+1)
log2(sRPKM+1)
PRC2cKO
C
Peaks
PRC1ctrl
PRC1cKO
PRC2ctrl
PRC2cKO
All H3K4me3 peaks (n=54,947)
All H3K4me3 peaks (n=56,326)
sRPKM
-3 center 3kb
D
Promoters
PRC1ctrl
PRC1cKO
PRC2ctrl
PRC2cKO
Genecode vm25 all genes (n=53,481)
Genecode vm25 all genes (n=53,481)
sRPKM
-2.5 TSS 2.5kb
E
Differential enrichment analyses
Padj<0.05;log2FoldChange>1
Lost in cKO
stable
Gained in cKO
Peaks
Promoters
PRC1ctrl vs. PRC1cKO
3.8%(n=1,970)
25.6%(n=13,159)
3.3%(n=1,747)
17.3%(n=9,261)
PRC2ctrl vs. PRC2cKO
13.6%(n=7,242)
4.9%(n=2,592)
6.0%(n=3,227)
4.5%(n=2,429)
F
PRC1ctrl vs. PRC1cKO Peaks
Lost in cKO (n=1,970)
3.08%
1.80%
69.43%
25.70%
Gained in cKO (n=13,159)
1.77%
2.34%
49.44%
46.45%
PRC2ctrl vs. PRC2cKO
Lost in cKO (n=7,242)
3.52%
1.86%
51.07%
43.55%
Gained in cKO (n=2,592)
1.72%
3.11%
71.78%
23.39%
TSS
TES
genebody
intergnenic
G
PRC1cKO H3K4me3 gained promoters (n=9,261)
ChEA
SUZ12 18555785 ChIP-Seq MESCs Mouse
RNF2 18974828 ChIP-Seq MESCs Mouse
EZH2 18974828 ChIP-Seq MESCs Mouse
JARID2 20075857 ChIP-Seq MESCs Mouse
EED 16625203 ChIP-ChIP MESCs Mouse
JARID2 20064375 ChIP-Seq MESCs Mouse
SUZ12 18974828 ChIP-Seq MESCs Mouse
SUZ12 18692474 ChIP-Seq MEFs Mouse
EZH2 27304074 Chip-Seq ESCs Mouse
SUZ12 18692474 ChIP-Seq MESCs Mouse
-logP

**Figure S9. H3K4me3 changes in the Polycomb conditional knockout mouse models.**
(A) Bar chart showing the foldchange of H3K4me3 global level in NGOs of PRC1ctrl&cKO and PRC2ctrl&cKO (n = 2 biological replicates, indicated by dots). *P* values of pairwise comparisons (two-sided unpaired Student's t-test) are given. Data are represented as mean ± SD.
(B) Density scatter plots showing genome-wide H3K4me3 enrichment by 10-kb bins, comparing PRC1ctrl&cKO, PRC2ctrl&cKO, respectively.
(C) Average tag density plots and heatmaps showing all H3K4me3 peaks in PRC1ctrl&cKO and PRC2ctrl&cKO NGOs.
(D) Average tag density plots and heatmaps showing H3K4me3 enrichment on all promoters in PRC1ctrl&cKO and PRC2ctrl&cKO NGOs.
(E) Bar charts showing percentages of differentially enriched peaks, and promoters for H3K4me3 in NGOs of PRC1ctrl&cKO, PRC2ctrl&cKO, respectively.
(F) Genomic distribution of diffenrential enriched H3K4me3 peaks in NGOs of PRC1ctrl&cKO, PRC2ctrl&cKO, respectively.
(G) Bar charts of ChIP-x Enrichment Analysis (ChEA) of gene promoters gaining H3K4me3 differentially in PRC1cKO NGOs. ChEA enrichment shows the transcription factors and the cell and animal types used in the profiling experiments.

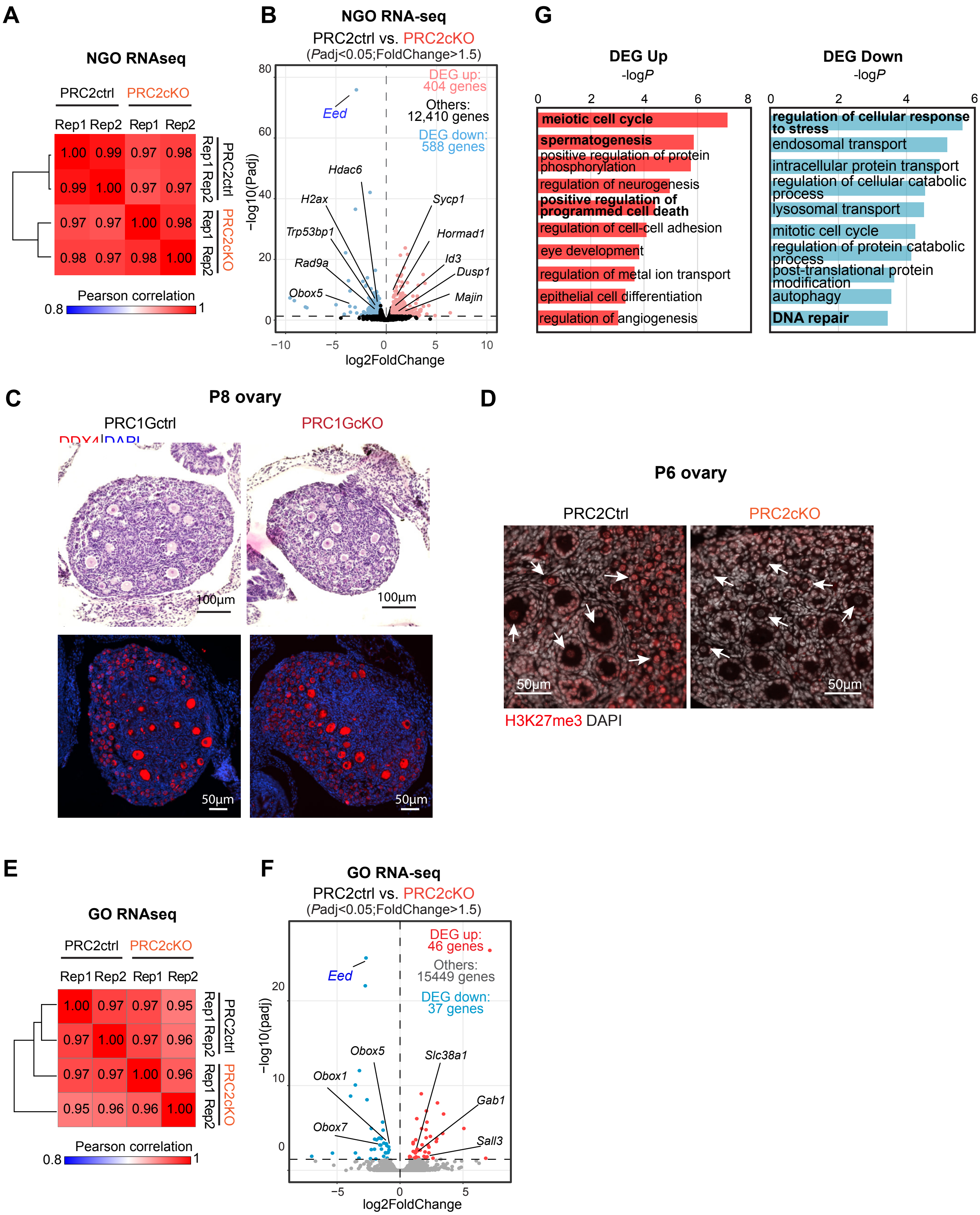

A
NGO RNAseq
PRC2ctrl
PRC2cKO
Rep1 Rep2 Rep1 Rep2
1.00 0.99 0.97 0.98
0.99 1.00 0.97 0.97
0.97 0.97 1.00 0.98
0.98 0.97 0.98 1.00
Pearson correlation
0.8
1
B
NGO RNA-seq
PRC2ctrl vs. PRC2cKO
(Padj<0.05;FoldChange>1.5)
DEG up: 404 genes
Others: 12,410 genes
DEG down: 588 genes
Eed
Hdac6
H2ax
Trp53bp1
Rad9a
Obox5
Sycp1
Hormad1
Id3
Dusp1
Majin
−log10(Padj)
log2FoldChange
G
DEG Up
-logP
meiotic cell cycle
spermatogenesis
positive regulation of protein phosphorylation
regulation of neurogenesis
positive regulation of programmed cell death
regulation of cell-cell adhesion
eye development
regulation of metal ion transport
epithelial cell differentiation
regulation of angiogenesis
DEG Down
-logP
regulation of cellular response to stress
endosomal transport
intracellular protein transport
regulation of cellular catabolic process
lysosomal transport
mitotic cell cycle
regulation of protein catabolic process
post-translational protein modification
autophagy
DNA repair
C
P8 ovary
PRC1Gctrl
PRC1GcKO
100µm
50µm
D
P6 ovary
PRC2Ctrl
PRC2cKO
50µm
H3K27me3 DAPI
E
GO RNAseq
PRC2ctrl
PRC2cKO
Rep1 Rep2 Rep1 Rep2
1.00 0.97 0.97 0.95
0.97 1.00 0.97 0.96
0.97 0.97 1.00 0.96
0.95 0.96 0.96 1.00
Pearson correlation
0.8
1
F
GO RNA-seq
PRC2ctrl vs. PRC2cKO
(Padj<0.05;FoldChange>1.5)
DEG up: 46 genes
Others: 15449 genes
DEG down: 37 genes
Eed
Obox5
Obox1
Obox7
Slc38a1
Gab1
Sall3
−log10(padj)
log2FoldChange

**Figure S10. Phenotypes of the Polycomb conditional knockout mouse models.**
(A, E) Heatmaps showing Pearson correlation values among each biological replicate in the PRC2 oocytes RNA-seq datasets.
(B, F) Volcano plots showing comparisons of transcriptomes between PRC2ctrl and PRC2cKO oocytes. All the genes with *P*adj values are plotted. Differentially expressed genes (DEGs: FoldChange >1.5, *P*adj< 0.05) are colored (red: up-regulated in PRC2cKO oocytes; blue: down-regulated in PRC2cKO oocytes), and numbers are shown.
(C) Histology of ovarian sections from PRC1ctrl and PRC1GcKO females at P8, stained with hematoxylin & eosin or immunostained for DDX4 (red). Two mice of each genotype were used for analysis, and representative images are shown.
(D) Immunostaining of H3K27me3 in ovaries of PRC2cKO and a control littermate at P6. H3K27me3 intensity is largely decreased in oocytes of PRC2cKO. Bars: 50 µm. White arrows point to the oocyte nucleus. At least three mice were analyzed for each genotype, and representative images are shown.
(G) Gene Ontology analyses of the DEGs in (B). Key Gene Ontology terms of the DEGs are shown. *P* values were generated by Metascape using the two-sided hypergeometric test.
